# The Hao-Fountain syndrome protein USP7 regulates neuronal connectivity in the brain via a novel p53-independent ubiquitin signaling pathway

**DOI:** 10.1101/2023.10.24.563880

**Authors:** Hao Chen, Cole J. Ferguson, Dylan C. Mitchell, Amanda Titus, Joao A. Paulo, Andrew Hwang, Tsen-Hsuan Lin, Hiroko Yano, Wei Gu, Sheng-Kwei Song, Carla M. Yuede, Steven P. Gygi, Azad Bonni, Albert H. Kim

## Abstract

Precise control of protein ubiquitination is essential for brain development, and hence, disruption of ubiquitin signaling networks can lead to neurological disorders. Mutations of the deubiquitinase USP7 cause the Hao-Fountain syndrome (HAFOUS), characterized by developmental delay, intellectual disability, autism, and aggressive behavior. Here, we report that conditional deletion of USP7 in excitatory neurons in the mouse forebrain triggers diverse phenotypes including sensorimotor deficits, learning and memory impairment, and aggressive behavior, resembling clinical features of HAFOUS. USP7 deletion induces neuronal apoptosis in a manner dependent of the tumor suppressor p53. However, most behavioral abnormalities in USP7 conditional mice persist despite p53 loss. Strikingly, USP7 deletion in the brain perturbs the synaptic proteome and dendritic spine morphogenesis independently of p53. Integrated proteomics analysis reveals that the neuronal USP7 interactome is enriched for proteins implicated in neurodevelopmental disorders and specifically identifies the RNA splicing factor Ppil4 as a novel neuronal substrate of USP7. Knockdown of Ppil4 in cortical neurons impairs dendritic spine morphogenesis, phenocopying the effect of USP7 loss on dendritic spines. These findings reveal a novel USP7-Ppil4 ubiquitin signaling link that regulates neuronal connectivity in the developing brain, with implications for our understanding of the pathogenesis of HAFOUS and other neurodevelopmental disorders.

## Introduction

As a fundamental post-translational modification, protein ubiquitination plays critical roles in neuronal development and function, and deregulation of ubiquitination is thought to contribute to the pathogenesis of diverse neurological conditions including neurogenetic disorders. For instance, perturbation of ubiquitin signaling has been implicated in distinct genetic disorders of brain development including Angelman^1^, Gordon-Holmes^2,3^ and Ferguson-Bonni^4^ syndromes. Substrate ubiquitination is regulated by both ubiquitin ligases, which catalyze the covalent attachment of the 76 amino acid-small ubiquitin protein to specific substrates, and deubiquitinases, which catalyze the removal of ubiquitin from specific substrates. Although ubiquitin ligases have been extensively studied for their roles in neurodevelopment^5^, the functions of most deubiquitinases in the nervous system have remained largely unexplored^6^, despite their link with neurodevelopmental disorders in genetic association studies^7^. We herein characterized the role of the deubiquitinase ubiquitin specific protease 7 (USP7) in the mammalian brain. Deletion or mutation of *USP7* causes the Hao-Fountain syndrome, which is characterized by developmental delay, intellectual disability, autism, and aggressive behavior^8,9^. However, the neuropathogenesis of the Hao-Fountain syndrome has remained poorly understood.

Studies in cancer biology have revealed that USP7 induces the turnover of the tumor suppressor protein p53^10,11^, and inhibition of USP7 may hold therapeutic promise for suppressing tumor growth^12,13^. A key target of USP7 is the ubiquitin ligase Mdm2, which ubiquitinates p53 and thereby promotes p53 degradation. USP7 counteracts the effects of Mdm2 auto-ubiquitination to stabilize Mdm2 and promote Mdm2-dependent ubiquitination and consequent degradation of p53^11^. In mice, global mutation of *Usp7* leads to early embryonic lethality, indicating the essential role of USP7 in embryogenesis^14^. Loss of USP7 in neural stem cells leads to smaller brain size, perinatal lethality, and p53 accumulation^15^. Although mutation of *Trp53*, the gene encoding p53 in mice, rescues brain size in *Usp7* mutant mice, co-deletion of p53 fails to improve newborn survival, indicating that USP7 plays essential roles in the nervous system through unknown p53-independent mechanisms^15^. Importantly, whereas USP7 function in proliferating cells has been the subject of intense scrutiny^16–18^, the roles and mechanisms of USP7 in post-mitotic neurons that might be relevant to the pathogenesis of the Hao-Fountain syndrome have remained largely unknown.

In this study, we conditionally knock out *Usp7* in post-mitotic glutamatergic neurons of the forebrain in mice. We find that USP7 mutant mice display distinct behavioral deficits that recapitulate many clinical features of the Hao-Fountain syndrome. We also find that USP7 loss induces neuronal apoptosis in the cerebral cortex during brain development in a p53-dependent manner. However, genetic deletion of p53 fails to rescue most behavioral deficits in USP7 mutant mice, suggesting that USP7 regulates brain development via p53-independent mechanisms. Accordingly, proteomic and dendritic spine morphological analyses reveal marked reduction in synapse formation in USP7 mutant mice independently of p53. An unbiased protein interaction and quantitative TMT-proteomic approach reveals, for the first time, the landscape of potential USP7 substrates in neurons, including the the SWI/SNF chromatin remodeling and spliceosomal complexes, implicating USP7 in fundamental aspects of chromatin and gene regulation in neurons. Specifically, we identify the RNA splicing factor Ppil4 as a novel functional substrate of USP7. Importantly, knockdown of Ppil4 recapitulates the dendritic spine phenotype observed in USP7 mutant mice. These findings define a novel ubiquitin signaling pathway, which links the deubiquitinase USP7 with the RNA splicing factor Ppil4 and drives neuronal connectivity in the developing brain, shedding light on the roles and mechanisms of USP7 in post-mitotic neurons, with important implications for our understanding of the neuropathogenesis of the Hao-Fountain syndrome.

## Results

### Conditional deletion of USP7 in glutamatergic excitatory neurons alters mouse behavior

To characterize the role of USP7 in post-mitotic glutamatergic neurons of the forebrain, we crossed mice harboring a floxed allele of *Usp7* (*Usp7*^fl/fl^) with mice expressing the recombinase Cre under the control of the regulatory sequence of the mouse *NEX* gene (*NEX-cre*)^19^ (Figure 1A). Immunofluorescence (IF) analyses demonstrated loss of USP7 in NeuN-positive cortical neurons of *Usp7*^fl/fl^; *NEX-cre* mice (hereafter *Usp7* cKO) (Figure 1B). USP7 loss occurred in post-mitotic neurons of the cortical plate but not in the ventricular zone, confirming that USP7 is specifically deleted in post-mitotic neurons (Figure S1A).

**Figure 1.**
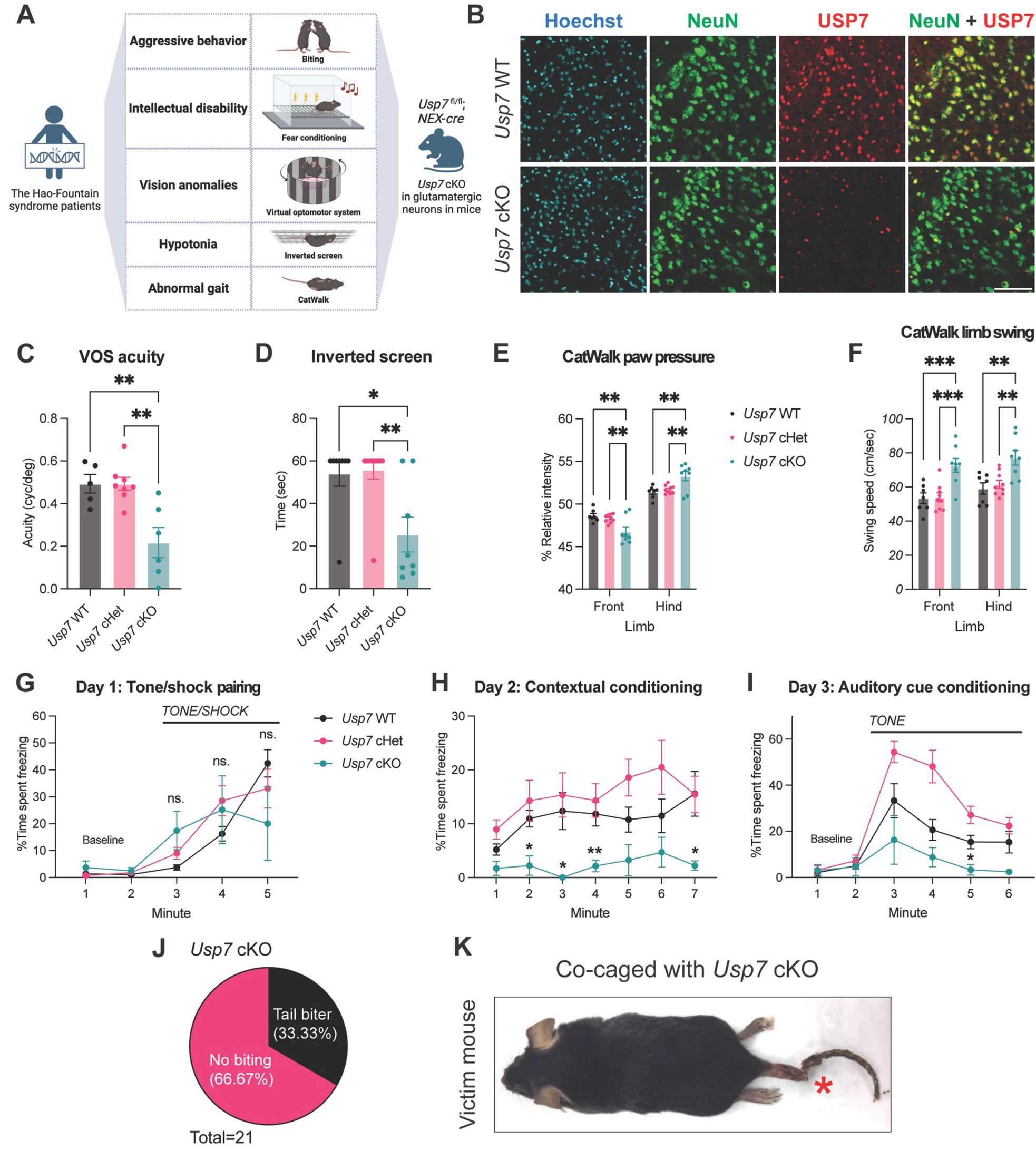
Loss of USP7 in glutamatergic excitatory neurons leads to multiple disease-relevant behavioral deficits in mice. (A) Patients with the Hao-Fountain syndrome and *Usp7* conditional knockout (cKO) mice exhibit phenotypes in similar behavioral domains. (B) Double immunofluorescence of USP7 and neuron nuclear protein (NeuN) in cerebral cortex at postnatal day 42 (P42). USP7 protein was absent in most neuronal nuclei in *Usp7* cKO cortex. Scale bar 100µm. (C) Virtual optomotor system (VOS) test: Spatial frequency threshold for head turn. *Usp7* cKO mice (n = 6) showed reduced visual acuity compared to *Usp7* WT (n = 5) and cHet mice (n = 8). **p<0.01 by Bonferroni’s multiple comparison test. (D) Motor strength test: Time to grip onto inverted screen. *Usp7* cKO mice (n = 8) showed shorter latency to fall than *Usp7* WT (n = 8) and cHet mice (n = 11). *p<0.05, **p<0.01 by Dunn’s multiple comparison test. (E) Catwalk 1: Relative mean intensity distributed onto each paw. *Usp7* cKO mice (n = 8) put less weight onto front paw and more onto hind paw, compared to *Usp7* WT (n = 7) and cHet mice (n = 9). **p<0.01 by Tukey’s multiple comparison test. See also Video S2. (F) Catwalk 2: Limb swing speed. *Usp7* cKO mice (n = 8) showed higher swing speed of both front and hind limb than *Usp7* WT (n = 7) and cHet mice (n = 9). **p<0.01, ***p<0.001 by Tukey’s multiple comparison test. See also Video S2. (G) Fear conditioning day 1: Time spent in freezing for tone/shock pairing (n = 8, 11 & 6 for *Usp7* WT, *Usp7* cHet and *Usp7* cKO mice). No significant genotype effect by two-way ANOVA. (H) Fear conditioning day 2: Time spent in freezing for contextual conditioning. *Usp7* cKO mice spent less time in freezing than *Usp7* WT mice. Numbers of mice are the same as in (G). *p<0.05, **p<0.01 by Bonferroni’s multiple comparison test (*Usp7* WT vs. cKO). (I) Fear conditioning day 3: Time spent in freezing for auditory cue conditioning. *Usp7* cKO mice spent less time in freezing after the tone than *Usp7* WT mice. Numbers of mice are the same as in (G). *p<0.05 by Bonferroni’s multiple comparison test (*Usp7* WT vs. cKO). (J) Percentage of tail biters among all monitored *Usp7* cKO mice from 3 to 12 weeks old. Tail biters were identified if their co-caged mice had biting marks on the tails. (K) Representative biting injury on mice co-caged with *Usp7* cKO mice. See also Video S3. Data are presented as mean ± SEM. See also Figure S1.

Patients with *USP7* mutation or deletion exhibit a range of deficits early in life that include growth retardation, hypotonia, visual impairment, gait abnormalities, aggression, and autism with intellectual disability^9^ (Figure 1A). To ascertain the effects of USP7 deletion on the organism level, we subjected *Usp7* cKO (*Usp7*^fl/fl^; *NEX-cre*), *Usp7* cHet (*Usp7*^fl/+^; *NEX-cre*), and *Usp7* WT (*Usp7*^fl/fl^) mice to a battery of behavioral tests and biometric measurements (Figure 1C-K and S1D-K). Although *Usp7* cKO mice exhibited small size during the juvenile period, the body weight of adult *Usp7* cKO mice was similar to controls (Figure S1B and S1C). This pattern of temporary growth delay is commonly seen in genetic mouse models of neurodevelopmental disorders^20^ and indicates that loss of USP7 in the brain is sufficient to generate phenotypes throughout the body.

Visual deficits are common in the Hao-Fountain syndrome^9^. To examine visual function in *Usp7* cKO mice, we employed the virtual optomotor system (VOS). In this experiment, mice make reflexive head movements to track the rotation of an alternating pattern of black and white vertical stripes, the spatial frequency of which can be experimentally varied. We found that *Usp7* cKO mice stopped responding at a much lower spatial frequency, suggesting reduced visual acuity (Figure 1C). This result shows that USP7 depletion in glutamatergic neurons is sufficient to compromise vision in mice.

Hypotonia is a motor impairment associated with the Hao-Fountain syndrome^8,9^. To assess motor performance in *Usp7* cKO mice, we began by performing the inverted screen test. In this strength test, *Usp7* cKO mice exhibited a significant reduction in the latency to fall (Figure 1D). We evaluated motor coordination using the ledge, platform, and pole tests. *Usp7* cKO mice also displayed a reduction in the latency to fall in the ledge and platform tests (Figure S1D and S1E) and were slower to turn around on the pole. (Figure S1G). However, mice of all genotypes spent similar amounts of time climbing down the pole (Figure S1F). These observations indicate that USP7 loss in the forebrain impairs motor strength and coordination without effects on simple motor tasks.

In the course of characterizing *Usp7* cKO mice, we noticed that they exhibited obvious gait abnormalities, another feature of USP7 mutation/deletion in humans^9^. During voluntary locomotion, *Usp7* cKO mice exhibited abnormal splaying of the hindlimbs (Video S1). To systematically evaluate this phenotype in *Usp7* cKO mice, we employed the CatWalk XT system to deeply analyze gait and detect subtle abnormalities (Video S2). These experiments demonstrated that *Usp7* cKO mice place extra weight onto their hind paws (Figure 1E). This analysis also showed an unexpected increase in the swing speed of the limbs of *Usp7* cKO mice (Figure 1F, S1H and S1I). Together, these results demonstrate that USP7 depletion in glutamatergic neurons of the forebrain leads to diverse but specific motor deficits in mice that mimic the motor phenotypes associated with the Hao-Fountain syndrome.

Having characterized the effect of conditional USP7 knockout on sensorimotor function, we next evaluated whether *Usp7* cKO mice exhibit deficits in tests of learning and memory. To avoid the potential confounding effect of visual impairment in *Usp7* cKO mice, we chose the fear conditioning paradigm using an auditory cue. We confirmed that hearing was not significantly impaired in *Usp7* cKO mice using the acoustic startle test (Figure S1J). On day 1, mice were exposed to an auditory tone (a cue) in a peppermint-scented chamber (the context) prior to the delivery of a mild electrical shock to the paw (Figure 1G). On day 2, during re-exposure to the context environment, *Usp7* cKO mice exhibited reduced freezing behavior (the conditioned response) (Figure 1H), suggesting diminished association between the context (the conditioned stimulus) and the shock (the unconditioned stimulus) in *Usp7* cKO mice. On day 3, upon re-exposure to the auditory cue (the conditioned stimulus) in a distinct context environment, *Usp7* cKO also exhibited diminished freezing behavior (the conditioned response) (Figure 1I). In contrast, in spite of their visual impairment, *Usp7* cKO mice showed no deficits in the Y-maze test of spatial working memory (Figure S1K). Collectively, these results demonstrate that USP7 depletion in glutamatergic neurons of the forebrain causes specific deficits in behavioral tests of associative learning.

Finally, *Usp7* cKO mice displayed striking forms of aggressive behavior characterized mainly by violent attacks on cage mates that resulted in wounds and occasionally amputation of the other animal’s tail (Video S3). Between the ages of 3 and 12 weeks, 7 out of 21 *Usp7* cKO mice inflicted bite marks centered around the tails of co-caged mice (Figure 1J and 1K). This behavior is highly unusual and distinct from the commonly observed biting that is associated with social aggression, which occurs after sexual maturity and typically targets the lower back and abdomen^21^.

In summary, this combination of observational and hypothesis-driven behavioral analyses demonstrates that loss of USP7 in glutamatergic neurons of the forebrain causes a suite of deficits that closely mimics the clinical hallmarks of the Hao-Fountain syndrome, including the unusual phenotype of aggression^8,9^. These results suggest that USP7 may play a conserved role in the brain in humans and mice. Therefore, the investigation of *Usp7* cKO mice should advance our understanding of ubiquitin signaling during brain development and the pathogenesis of neurodevelopmental disorders.

### USP7 prevents p53-dependent neuronal apoptosis

After determining that loss of USP7 in the forebrain triggers diverse behavioral deficits in mice, we sought to determine how USP7 regulates neuronal development in the cerebral cortex. Therefore, we performed histological analyses at different stages of neurodevelopment, beginning at embryonic day 12 (E12) to coincide with the onset of *NEX-Cre* expression^19^. The earliest phenotype that we detected was a marked increase in the number of cells that were positive for the apoptosis marker protein cleaved caspase 3 (c-cas 3) by immunofluorescence in the cerebral cortex in *Usp7* cKO mice (Figure 2A). Importantly, this phenotype was temporally restricted in the cerebral cortex of *Usp7* cKO mice to the postnatal day 0 (P0) timepoint, and little difference in c-cas 3 was present at P5, P10 or E15 in control and *Usp7* cKO mice. These results show that USP7 loss in cortical glutamatergic neurons leads to a dramatic wave of apoptosis during the perinatal period.

**Figure 2.**
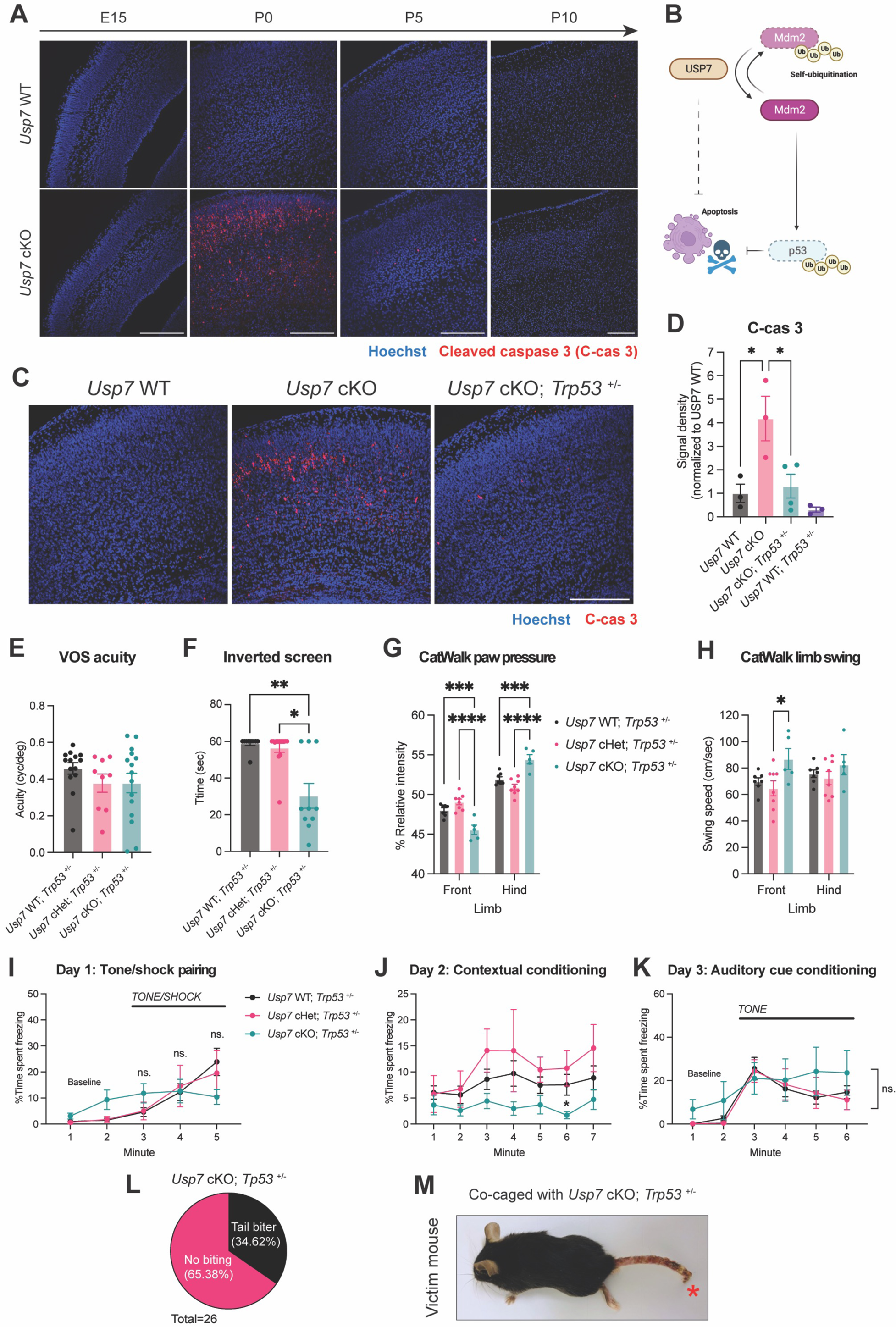
Deletion of p53 rescues apoptosis, impairment of visual acuity and auditory cue conditioning in *Usp7* cKO mice but not other behavioral deficits. (A) Immunofluorescence of apoptosis marker cleaved caspase 3 (c-cas 3) in cerebral cortex at different developmental time points. *Usp7* cKO cortex showed large scale of apoptosis at P0, whereas apoptosis is low at E15, P5 and P10. Scale bars all 200µm. (B) Schematic of USP7-Mdm2-p53 signaling pathway regulating apoptosis. (C) Immunofluorescence of c-cas 3 in P0 cortex of different *Trp53* and *Usp7* genotypes. *Trp53* heterozygosity diminished USP7 loss-induced apoptosis. Scale bar 200µm. (D) Quantification of c-cas 3 fluorescent signal, as shown in (C). Number of mice = 3, 3, 4 & 3 for *Usp7* WT, *Usp7* cKO, *Usp7* cKO; *Trp53* ^+/−^ & *Usp7* WT; *Trp53* ^+/−^, respectively. *p<0.05 by Tukey’s multiple comparison test. (E) Virtual optomotor system (VOS) test: Spatial frequency threshold for head turn. *Usp7* cKO; *Trp53*^+/−^ mice (n = 15) did not have impaired visual acuity, compared to *Usp7* WT; *Trp53* ^+/−^ (n = 14) and *Usp7* cHet; *Trp53* ^+/−^ mice (n = 9). No significant genotype effect by one-way ANOVA. (F) Motor strength test: Time to grip onto inverted screen. *Usp7* cKO mice (n = 10) showed shorter latency to fall than *Usp7* WT; *Trp53*^+/−^ (n = 10) and *Usp7* cHet; *Trp53*^+/−^ mice (n = 14). *p<0.05, **p<0.01 by Dunn’s multiple comparison test. (G) Catwalk 1: Relative mean intensity distributed onto each paw. *Usp7* cKO; *Trp53*^+/−^ mice (n = 5) put less weight onto front paw and more onto hind paw than *Usp7* WT; *Trp53*^+/−^ (n = 7) and *Usp7* cHet; *Trp53*^+/−^ mice (n = 8). ***p<0.001, ****p<0.0001 by Tukey’s multiple comparison test. See also Video S4. (H) Catwalk 2: Limb swing speed. *Usp7* cKO; *Trp53*^+/−^ mice showed slightly higher swing speed than *Usp7* cHet; *Trp53*^+/−^ mice. Numbers of mice are the same as in (G). *p<0.05 by Tukey’s multiple comparison test. See also Video S4. (I) Fear conditioning day 1: Time spent in freezing for tone/shock pairing (n = 12, 6 & 11 for *Usp7* WT, *Usp7* cHet and *Usp7* cKO mice). No significant genotype effect by two-way ANOVA. (J) Fear conditioning day 2: Time spent in freezing for contextual conditioning. *Usp7* cKO; *Trp53*^+/−^ mice spent less time in freezing than *Usp7* WT; *Trp53*^+/−^ mice. Numbers of mice are the same as in (H). *p<0.05, by Bonferroni’s multiple comparison test (*Usp7* WT vs. cKO). (K) Fear conditioning day 3: Time spent in freezing for auditory cue conditioning. *Usp7* cKO; *Trp53*^+/−^ mice had no difference in freezing after the tone, compared to *Usp7* WT; *Trp53*^+/−^ and *Usp7* cHet; *Trp53*^+/−^ mice. Numbers of mice are the same as in (H). No significant genotype effect by two-way ANOVA. (L) Percentage of tail biters among all monitored *Usp7* cKO; *Trp53*^+/−^ mice from 3 to 12 weeks old. Tail biters were counted if their co-caged mice had biting marks on their tails. (M) Representative biting injury on mice co-caged with *Usp7* cKO; *Trp53*^+/−^ mice. Data are presented as mean ± SEM. See also Figure S2, S3 and S4.

To determine whether neuronal apoptosis in the cerebral cortex of *Usp7* cKO mice is cell-autonomous, we used immunofluorescence to simultaneously detect Cre and c-cas 3. This analysis demonstrated considerable overlap between Cre and c-cas 3 (Figure S2A), suggesting that apoptosis occurs specifically in Cre-expressing glutamatergic neurons. In other experiments, RNA interference (RNAi)-induced knockdown of USP7 sparsely in primary cortical neurons using calcium phosphate transfection robustly induced apoptosis (Figure S2B-D). Together, these results suggest that USP7 loss triggers neuronal apoptosis in a cell-autonomous manner.

Because USP7 regulates the Mdm2-p53 pathway in cancer and proliferating cells^9,10,14^ (Figure 2B), we asked whether the apoptosis in post-mitotic neurons upon USP7 knockout in the mouse cerebral cortex is mediated by p53. We crossed the *Usp7* cKO mice with mice harboring a null allele in *Trp53*, the gene that encodes p53 in mice. Notably, the loss of just one *Trp53* allele in *Usp7* cKO mice reduced the c-cas 3 signal to a level that was comparable to control mice (Figure 2C and 2D). Homozygous knockout of *Trp53* rescued apoptosis in the cortex of *Usp7* cKO mice to a comparable degree as for heterozygous loss of p53 (Figure S3A). Together, these results demonstrate that p53 is required for neuronal apoptosis in *Usp7* cKO animals (Diagram as Figure 2B).

To test whether the rescue of apoptosis in *Usp7* cKO neurons by p53 loss is cell-autonomous, we sought to assess the effect of p53 depletion on apoptosis in primary cultures of cortical neurons. We began by isolating neurons from E15 mouse embryos harboring floxed *Usp7* (*Usp7* ^fl/fl^). Following transduction with a lentivirus encoding Cre, immunocytochemical analyses confirmed loss of USP7 upon Cre transduction (Figure S3B). In contrast, neurons transduced with a catalytically dead Cre-Y331F mutant (Cre dead) failed to show depletion of USP7. In cortical neurons dissociated from *Usp7*^fl/fl^; *Trp53* ^+/+^ embryos, loss of USP7 following wild-type Cre (Cre WT) transduction substantially reduced cell viability in the MTS assay. In contrast, neurons from *Usp7* ^fl/fl^; *Trp53* ^+/−^ and *Usp7* ^fl/fl^; *Trp53* ^−/−^ embryos demonstrated no significant decrease in viability upon Cre WT expression (Figure S3C). These results suggest that p53 deletion rescues USP7-dependent cell death in a cell-autonomous manner.

Our observation that deletion of USP7 triggers neuronal apoptosis led us to ask whether this cellular phenotype might translate to a reduction in brain volume. Using diffusion-weighted *ex vivo* MRI, we found that the volumes of both the cerebral cortex and the hippocampal formation were dramatically reduced by ∼40% in *Usp7* cKO mice (Table 1, Figure S3D and S3E). Surprisingly, although p53 loss to rescued apoptosis in the brains of *Usp7* cKO mice, co-deletion of p53 in *Usp7*^fl/fl^; *NEX-cre*; *Trp53* ^+/−^ and *Usp7*^fl/fl^; *NEX-cre*; *Trp53* ^−/−^ mice had minimal effect on the volumes of the cerebral cortex or hippocampus (Table 1, Figure S3D and S3E). In agreement with these observations in the forebrain, total brain volume was reduced by ∼20% in both *Usp7*^fl/fl^; *NEX-cre*; *Trp53*^+/+^ and *Usp7* ^fl/fl^; *NEX-cre*; *Trp53*^+/−^ mice (Table 1, Figure S3D and S3E). These results suggest that p53-mediated apoptosis does not significantly contribute to the reduction of brain volume in USP7 cKO mice, indicating the importance of p53-independent pathways to this phenotype.

**Table 1.** Phenotype comparison between *Usp7* cKO mice and *Usp7* cKO; *Trp53*^+/−^ mice. Phenotypes significantly different from *Usp7* WT or *Usp7* WT; *Trp53*^+/−^ mice are in bold. The whole brain volumes exclude olfactory bulbs. Hippo = hippocampus. na = not available.

| Test | Phenotype | *Usp7* cKO mice | *Usp7* cKO; *Trp53*<sup>+/-</sup> mice |
| --- | --- | --- | --- |
| **c-cas 3 immunofluorescence** | **Apoptosis** | **Ectopically induced** | **Normal** |
| *Ex vivo* MRI | Brain volume | -37.4% (cortex)<br>-39.1% (hippo)<br>-20.0% (whole brain) | -26.3% (cortex)<br>-30.9% (hippo)<br>-21.3% (whole brain) |
| Body weight measurement | Body weight | Growth delay | Growth delay |
| **VOS test** | **Visual acuity** | **Impaired** | **Normal** |
| Inverted screen test | Motor strength | Impaired | Impaired |
| Ledge, platform and pole tests | Motor coordination | Impaired | Impaired |
| Catwalk | Gait | Impaired | Impaired |
| Acoustic startle response | Hearing ability | Normal | Normal |
| **Fear conditioning** | Contextual conditioning | Impaired | Impaired |
|  | **Cued conditioning** | **Impaired** | **Normal** |
| Y maze | Spatial working memory | Normal | na |
| Tail biting | Aggression | Observed in 33.3% mice | Observed in 34.62% mice |

### p53 reduction rescues a subset of behavioral abnormalities in *Usp7* cKO mice

To determine whether dysregulation of p53 contributes to behavioral deficits in *Usp7* cKO mice, we measured biometrics and tested motor performance and memory formation in mice of the genotype *Usp7*^fl/fl^; *NEX-cre*; *Trp53*^+/−^ (hereafter *Usp7* cKO; *Trp53*^+/−^) (Figure 2E-M and S4A-I). The growth of *Usp7* cKO; *Trp53*^+/−^ juvenile mice remained delayed compared to control mice of the genotypes *Usp7*^+/+^; *NEX-cre*; *Trp53*^+/−^ (hereafter *Usp7* WT; *Trp53*^+/−^) and *Usp7*^fl/+^; *NEX-cre*; *Trp53*^+/−^ (hereafter *Usp7* cHet; *Trp53*^+/−^) (Figure S4A and S4B). These findings suggest that juvenile growth delay that occurs upon USP7 loss in the forebrain does not require p53 as an intermediary. In contrast, *Usp7* cKO; *Trp53*^+/−^ mice demonstrated comparable visual acuity to *Usp7* WT; *Trp53* ^+/−^ mice, suggesting that visual impairment observed in *Usp7* cKO mice requires p53 (Figure 2E). Assessment of motor function in *Usp7* cKO; *Trp53*^+/−^ mice demonstrated persistent impairment in the inverted screen test (Figure 2F), suggesting that motor weakness associated with USP7 depletion in the brain persists despite p53 depletion. Similarly, *Usp7* cKO; *Trp53*^+/−^ mice were also impaired in the ledge, platform, and pole tests (Figure S4C-F). p53 deletion also failed to normalize the performance of *Usp7* cKO mice in the CatWalk, as *Usp7* cKO; *Trp53*^+/−^ also improperly distributed their body weight and accelerated the swing speed of their limbs during voluntary locomotion (Figure 2G, 2H, S4G-H and Video S4). Tail biting behavior was also present in *Usp7* cKO; *Trp53*^+/−^ mice (Figure 2L and 2M). These results indicate that the motor impairment and aggression phenotypes in *Usp7* cKO mice are largely independent of p53 loss.

Finally, we analyzed the effect of p53 loss on the performance of *Usp7* cKO mice during associative learning paradigms. Similar to *Usp7* cKO mice, *Usp7* cKO; *Trp53*^+/−^ mice exhibited decreased freezing during contextual conditioning (Figure 2I and 2J). Prior to performing cued conditioning, we used the acoustic startle test to confirm that hearing was unimpaired in *Usp7* cKO; *Trp53*^+/−^ mice (Figure S4I). During cued fear conditioning, the impairment in the freezing response to the auditory cue in *Usp7* cKO mice was corrected in mice of the genotype *Usp7* cKO; *Trp53*^+/−^ (Figure 2K). Altogether, our behavioral analyses demonstrate that whereas p53 contributes to USP7 loss-induced impairment of vision and cued conditioning, most behavioral deficits in *Usp7* cKO mice are independent of p53 reduction and its effects on apoptosis (Table 1).

### Loss of USP7 reduces synapses in cortical neurons

After observing that most developmental phenotypes associated with USP7 mutation did not require p53, we attempted to identify USP7-dependent phenotypes in neurons beyond p53-mediated apoptosis. To probe cellular and molecular abnormalities upon USP7 depletion in the developing brain, we profiled the proteome of the cerebral cortex of *Usp7* cKO mice using unbiased tandem mass tag-mass spectrometry (TMT-MS)-based proteomic methods (Figure 3A). We included cortices from 5 pairs of sex-matched *Usp7* cKO mice and their control littermates at age P17. Protein lysates were trypsinized and labeled with individual tandem mass tag prior to quantitation of relative protein abundance using MS3. We sorted the differentially abundant proteins based on fold change and p-value in the unpaired t-test (Figure 3B). Although more proteins were up-regulated than down-regulated by loss of USP7, gene set enrichment analysis (GSEA) showed the most significantly enriched gene sets were among the down-regulated proteins (Figure 3C and Table S1). Importantly, synapse-related gene sets were prominent among the enriched gene sets (Figure 3C and Table S1). Additional GO analysis using the GOrilla algorithm^22^ also identified synapse-related terms among the ranked list of proteins whose abundance was reduced by USP7 loss (fold change < 0.75) (Figure 3D). Further SynGO analysis^66^ revealed that both postsynapse and presynapse terms are enriched in the down-regulated proteins by USP7 loss (Figure S5A&B). The down-regulated presynaptic proteins include Cplx3, Kcnj11, Il1rapl2 and Rpl35, and down-regulated postsynaptic proteins include Arc, Rtn4, Rnf220 and Dclk1 (Figure S5C). These results suggest that USP7 may regulate synaptic proteins in the cerebral cortex.

**Figure 3.**
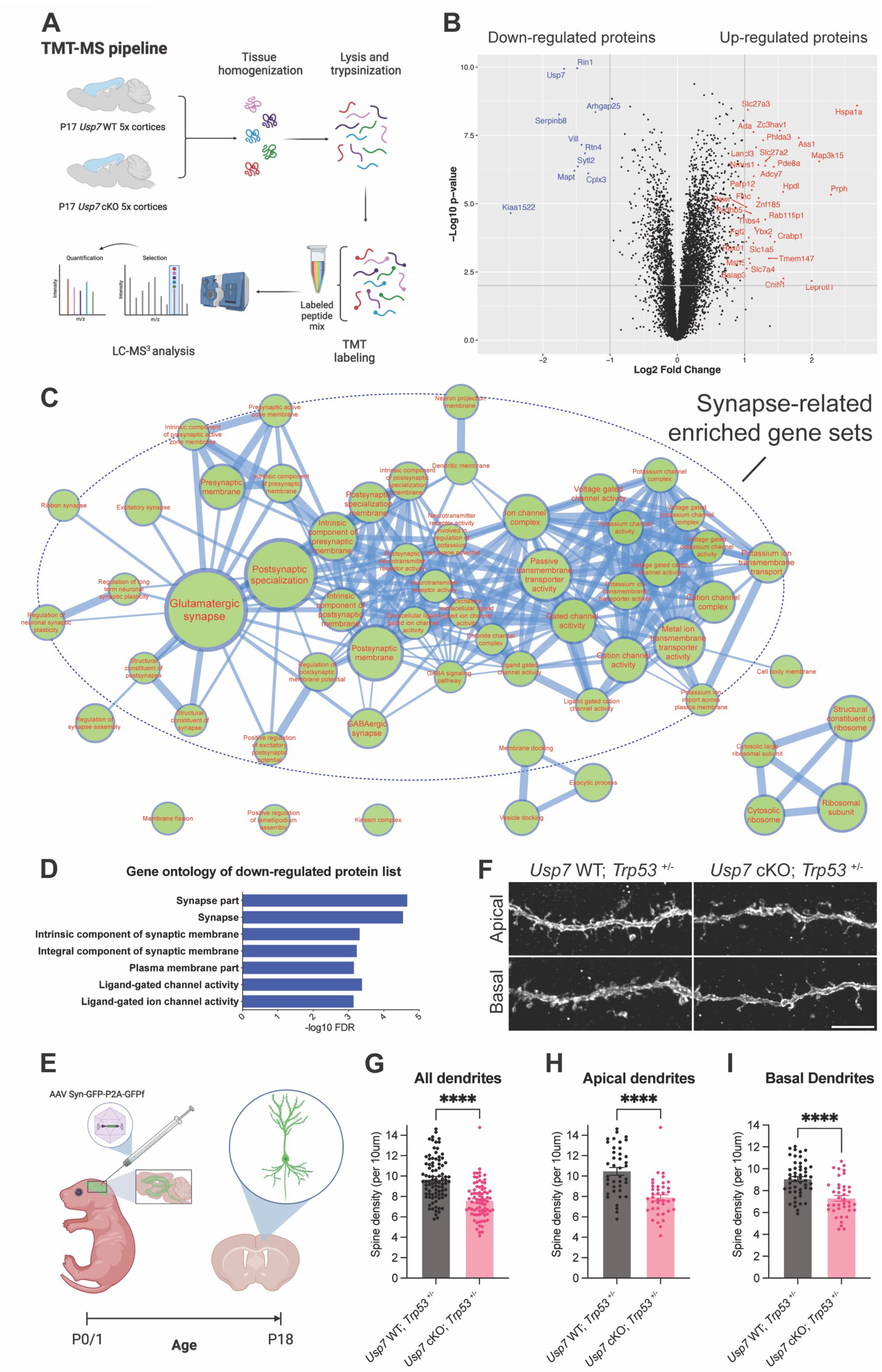
Loss of USP7 reduces synapses in cortex. (A) Flowchart to profile the proteome of the cerebral cortex of *Usp7* cKO mice with tandem mass tag-mass spectrometry (TMT-MS). (B) Volcano plot of all profiled proteins in TMT-MS. Up-regulated and down-regulated proteins (*Usp7* cKO vs. WT) are labeled in red and blue, respectively. P-value were calculated by 2-tail unpaired t-test. (C) Organic layout of enriched GO gene sets based on TMT protein abundance in (B). Nodes represent gene sets (q-value < 0.01) and size of nodes represents how many genes contained in gene sets. Edges represent GO-defined relations (similarity > 0.5) and thickness of edges represents similarity between gene sets. Most enriched gene sets are related to synapse and circled in red. All gene sets shown are enriched in *Usp7* WT and de-enriched in *Usp7* cKO. (D) GO terms enriched from ranked list of down-regulated proteins (*Usp7* cKO vs. WT fold change < 0.75). (E) Ventricular injection of AAV PHP.eB Syn-GFP-P2A-GFPf at P0/1 to sparsely label pyramidal neurons in cerebral cortex and dendritic spine analysis at P18. (F) Representative GFP micrographs of dendritic spines on apical and basal dendrites. Scale bar 5µm (G, H, I) Spine density on apical, basal and all dendrites of pyramidal neurons in motor cortex. *Usp7* cKO; *Trp53*^+/−^ neurons (n = 39 & 39 neurons for apical dendrites & basal dendrites from 3 mice) showed reduced spine density than *Usp7* WT; *Trp53*^+/−^ neurons (n = 40 & 50 neurons for apical dendrites & basal dendrites from 3 mice). ****p<0.0001 by 2-tail unpaired t-test Data are presented as mean ± SEM. See also Figure S5&6.

To test whether USP7 loss impairs synapse development, we next characterized the effects of USP7 loss on dendritic spines in glutamatergic neurons *in vivo*. We employed adeno-associated virus (AAV)-PHP.eB harboring a syn-GFP-P2A-GFPf construct to drive expression of green fluorescent protein (GFP) in neurons directed by the synapsin promoter. AAV was injected into the ventricles of newborn *Usp7* WT and *Usp7* cKO mouse pups at titers that produced sparse labeling of neurons, highlighting their dendritic spine morphology with cytoplasmic GFP and membrane bound GFPf (Diagram as Figure 3E). To avoid the influence of p53-mediated apoptosis in neurons, we analyzed the morphologic effects of USP7 loss on the *Trp53*^+/−^ background. Spine density within both apical and basal dendrites of pyramidal neurons in the motor cortex of *Usp7* cKO; *Trp53*^+/−^ mice at P18 was significantly reduced compared to that of *Usp7* WT; *Trp53*^+/−^ mice (Figure 3F-I). In spine shape analyses, filopodial spines were increased in *Usp7* cKO; *Trp53*^+/−^ neurons as compared to *Usp7* cKO; *Trp53*^+/−^ neurons (Figure S5D-J). We also found that dendrite length was unaffected in Cre-transduced *Usp7*^fl/fl^; *Trp53*^+/−^ primary cortical neurons, and there was no effect on dendrite arborization by Sholl analysis upon USP7 loss (Figure S6A-F). These results demonstrate that the loss of USP7 specifically reduces dendritic spines of pyramidal neurons in the developing mouse brain.

### Defining the interactome and candidate substrates of USP7 in neurons

Having identified critical p53-independent roles for USP7 in regulating dendritic spines in the developing mouse cerebral cortex, we next characterized the mechanism by which USP7 controls synapse formation in post-mitotic neurons. Because USP7 substrates in cancer cells frequently bind to the TRAF domain of USP7^18,23^ (Figure 4A), we reasoned that neuronal USP7 substrates might also interact with the TRAF domain of USP7 and thus characterized the interactors of the TRAF domain in primary cortical neurons. In co-immunoprecipitation (co-IP) analyses, we confirmed that the truncated USP7 TRAF domain interacts with the known USP7 substrate Mdm2. Alanine substitution mutations at key residues of the TRAF domain including Arg104 (R104A), Asp165 (D165A) and Trp166 (W166A), collectively termed the RDW mutant, disrupted the interaction between the TRAF domain and Mdm2 (Figure S7A). Immunocytochemical analyses in primary cortical neurons confirmed the mostly nuclear localization of both the wild-type TRAF domain and the RDW mutant TRAF domain (Figure S6B and S7C). Silver staining of lysates from primary cortical neurons expressing lentiviral FLAG-tagged TRAF domain showed a large number of proteins that co-immunoprecipitated with the wild-type TRAF domain. Most of these putative interactions were lost by RDW mutations of the TRAF domain. (Figure 4B). Using the RDW mutant as a negative control, we performed mass spectrometry analysis of interactors of the USP7-TRAF domain (IP-MS) in 3 biological replicates of primary cortical neurons. We ranked interactors with the wild-type TRAF domain in neurons according to protein length-normalized spectral counts and generated an association network of the top 50 interactors based on the STRING (Search Tool for the Retrieval of Interacting Genes/Proteins) database. GO terms related to cellular components revealed a few densely interconnected structures on the STRING network that included ribonucleoprotein complexes, the RNA spliceosome, and the SWI/SNF (BAF) chromatin remodeling complex (Figure 4C and 4D). These results indicate that USP7 might interact with and/or target substrates involved in critical processes related to chromatin and RNA splicing in a TRAF domain-dependent manner.

**Figure 4.**
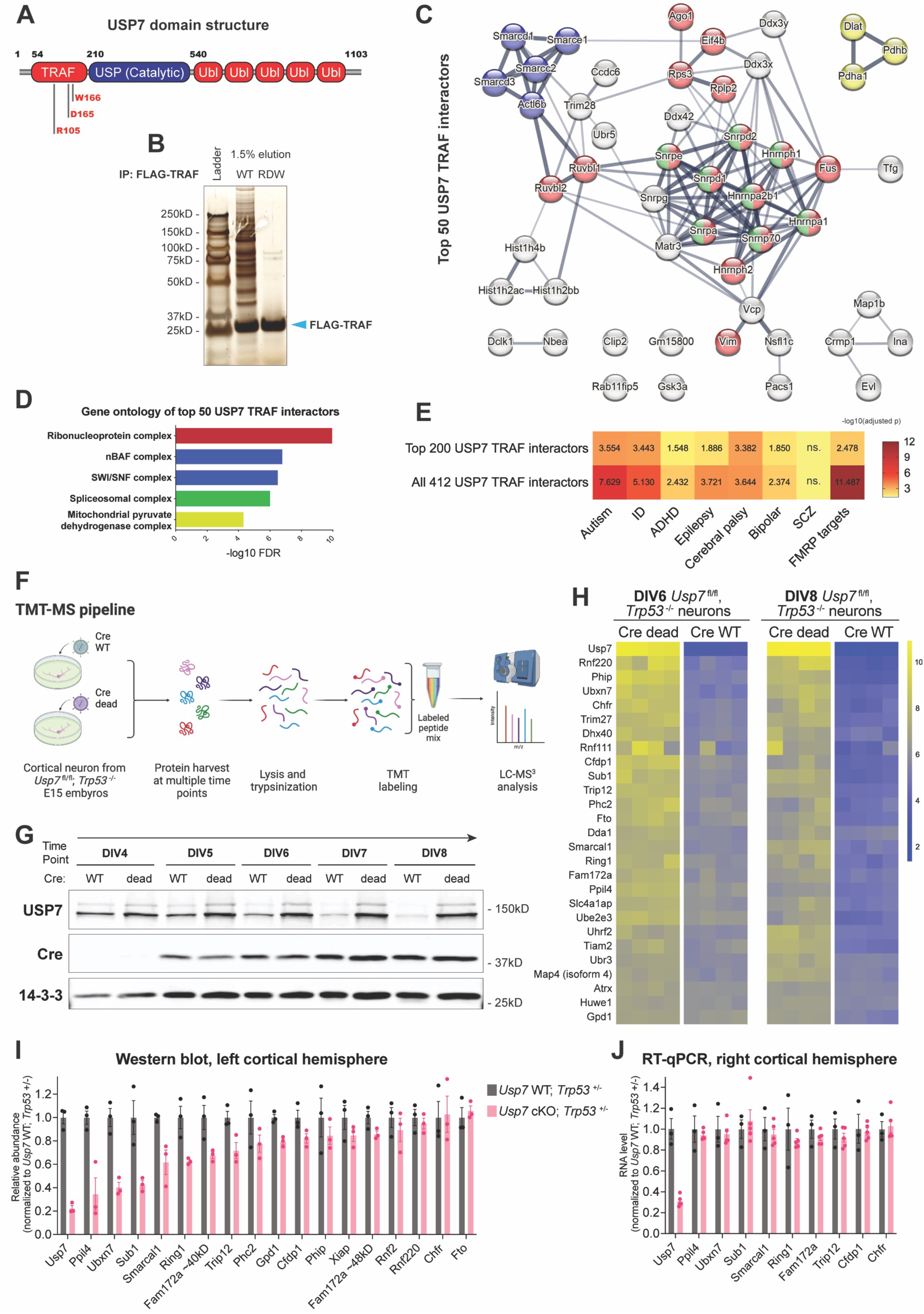
Unbiased proteomic characterizations of USP7 in cortical neurons. (A) Domain structure of mouse USP7 protein. R105, D165 and W165 on the TRAF domain are three amino acids critical for substrate binding^18,23^. USP: Ubiquitin specific protease domain. Ubl: Ubiquitin like domain. (B) Silver stain of FLAG-immunoprecipitated USP7 TRAF domain (arrow) with and without RDW mutations (R105A + D165A + W166A).1.5% of all eluted sample were stained while rest of the sample were analyzed by liquid chromatography-mass spectrometry (LC-MS). (C) Protein-protein association map of USP7 TRAF interactors (full STRING network, confidence > 0.4). Edge thickness represents confidence in association between proteins. Shown are only 50 interactors with highest gene length-normalized spectral counts. Ribonucleoprotein complex is colored in red. Spliceosomal complex is colored in green. SWI/SNF complex is colored in blue. Mitochondrial pyruvate dehydrogenase complex is colored in yellow. (D) Molecular component GO enrichments of top 50 interactors of the USP7 TRAF domain. Different molecular components are colored as in (C). (E) Disease gene enrichment analysis of the interactors of USP7 TRAF domain. ID = intellectual disability. SCZ = schizophrenia. ns. = not significant. (F) Flowchart to profile proteomic change with tandem mass tag-mass spectrometry (TMT-MS) in response to acute *Usp7* knockout in primary cortical neurons. Neurons were dissociated from *Usp7*^fl/fl^; *Trp53*^−/−^ embryos at E15 and subject to Cre lentiviral transduction at DIV3. Cre harboring Y331F mutation is catalytically dead (Cre dead) and served as negative control. (G) Western blot validation of USP7 depletion and Cre expression in Cre WT-transduced neurons over time. 14-3-3 is loading control. (H) Heatmap of TMT protein abundance of USP7 and its candidate substrates. Candidate substrates were sorted out by smallest p-value of Cre main effect (two-way ANOVA) and shortest Euclidean distance to USP7. (I) Western blot quantitation of candidate substrates with the cerebral cortex of *Usp7* cKO; *Trp53*^+/−^ (n = 3 mice) vs. *Usp7* WT; *Trp53*^+/−^ (n = 3 mice) at P0. Western blot images are in Figure S9. (J) RT-qPCR of candidate substrates with the cerebral cortex of *Usp7* cKO; *Trp53*^+/−^ (n = 5 mice) vs. *Usp7* WT; *Trp53*^+/−^ (n = 3 mice) at P0. No candidate substrate in the analysis showed RNA reduction with the loss of USP7. Data are presented as mean ± SEM. See also Figure S7, S8 and S9.

Among interactors of wild-type USP7 TRAF domain, a number of proteins are genetically associated with neurodevelopmental diseases (Ddx3x, Nbea, Smarcc2 and Map1b). Therefore, we compared curated neurodevelopmental disease gene sets^24^ with interactors of the USP7 TRAF domain. We found that USP7 TRAF interactors are most significantly enriched in the gene sets for intellectual disability and autism (Figure 4E), which are also the most common comorbilities in the Hao-Fountain syndrome^9^. Interestingly, USP7 TRAF interactors also had strong enrichment in the high-confidence mRNA targets of Fragile X syndrome protein FMRP^68^ (Figure 4E). These observations suggest that USP7 may interact with and target proteins implicated in neurodevelopmental disorders.

USP7 stabilizes its substrates by removing ubiquitin tags and preventing subsequent degradation. Therefore, we also searched for candidate USP7 substrate by profiling protein abundance using quantitative TMT-MS (Figure 4F). To minimize the influence of secondary effects that could occur in the setting of extended loss of USP7, we acutely inactivated USP7 by transducing primary cortical neurons isolated from mouse embryos of the genotype *Usp7*^fl/fl^; *Trp53* ^−/−^ with a lentivirus expressing the recombinase Cre. Immunoblotting analyses at different time points after transduction confirmed depletion of USP7 in lysates of primary cortical neurons (Figure 4G). Transduction and expression of catalytically dead Cre failed to induce USP7 loss (Figure 4G). Based upon the timing of USP7 depletion in this system, we selected DIV6 and DIV8 for TMT proteomics analysis. Consistent with the immunoblotting analyses, the TMT abundance of USP7 protein decreased at both DIV6 and DIV8 following transduction of wild-type Cre (Figure 4H). We identified and quantitated >8000 proteins, and principal component analysis (PCA) over these proteins showed clear segregation into the different experimental groups (Figure S8A). The first principal component accounted for ∼20% of the variance and separated samples at DIV6 from samples at DIV8. The second principal component accounted for ∼10% of the variance and separated the wild-type Cre versus dead Cre samples. Importantly, the distance between wild-type Cre versus dead Cre samples increased from DIV6 to DIV8, consistent with more complete depletion of USP7 at DIV8. These results demonstrate that acute USP7 loss induces a progressive proteomic change over time in primary cortical neurons.

Because USP7 stabilizes its substrates, we reasoned that the abundance of USP7 should correlate with the abundance of its substrates. Using this rationale to identify putative USP7 substrates, we calculated the Euclidian distance between USP7 and each protein detected in all 16 samples. We then analyzed the p-value of Cre-dependent effects for each protein using two-way ANOVA. Based on these analyses, we sorted out candidate substrates with the shortest distances to USP7 and the smallest p-values (Figure 4H). The abundance of each candidate substrate was down-regulated by expression of wild-type Cre at both DIV6 and DIV8. As proof of principle, this approach identified the known USP7 substrate Trim27^8^. Among proteins that were down-regulated by USP7 loss was the SWI/SNF subunit Smarcal1, the RNA spliceosome proteins Ppil4 and Fam172a, and several ubiquitin ligases including Trip12 and Ring1.

We next asked if the protein levels of these candidate substrates also decrease upon USP7 knockout *in vivo*. In our *in vivo* TMT analysis in the cerebral cortex of *Usp7* cKO mice, most of the candidate substrates that we found to be down-regulated upon USP7 loss in primary neurons also showed varying degrees of reduction in the cortex of P17 *Usp7* cKO mice (Figure S8B). To characterize these candidate substrates, we performed immunoblotting analysis of the P0 cererbal cortex of *Usp7* cKO; *Trp53* ^+/−^ mice. Performing these assays in the setting of *Trp53* depletion minimized the effects of apoptosis on our analysis. This analysis confirmed that several candidate substrates, including Ppil4, Ubxn7, Sub1, Smarcal1, Ring1, Fam172a and Trip12, were less abundant in the cortex of *Usp7* cKO; *Trp53*^+/−^ mice (Figure 4I and S9A-I). RT-qPCR showed no evidence that the abundance of any of the corresponding mRNAs was reduced by USP7 loss (Figure 4J), suggesting that USP7 regulates the abundances of these candidate substrates at the protein level. Altogether, these results demonstrate that USP7 regulates the abundance of specific candidate substrates in primary neurons and in the brain.

To better understand the global molecular network that is regulated by USP7, we used weighted correlation network analysis (WGCNA) to cluster proteins that become differentially abundant upon USP7 loss. We used a cutoff p-value of 0.05 for determining both Cre-dependent and Cre-time interaction effects using two-way ANOVA. WGCNA analysis identified 10 distinct network modules (Figure S8C). The yellow module, which contained most of the candidate substrates, showed down-regulation of its eigengene upon USP7 loss (Figure S8D and S8E). GO analysis of proteins in the yellow module revealed an enrichment in ubiquitin ligases (Figure S8F). Interestingly, ribonucleoprotein and splicing-related GO terms were overrepresented in the turquoise module, which consisted of the largest set of differentially abundant proteins among all the modules (Figure S8C and S8F). Collectively, findings from these unbiased analyses suggest that USP7 transforms the neuronal proteome by regulating a diverse network of proteins.

### USP7 targets Ppil4 to promote synaptogenesis

After profiling the interactome and putative targets of USP7 in neurons, we sought to further use these datasets to identify the biologically relevant USP7 targets in neurons. TMT proteomics followed by immunoblotting validation *in vivo* revealed that USP7 stabilizes Ppil4, Ubxn7, Sub1, Smarcal1, Ring1, Fam172a and Trip12, and IP-MS experiments identified a number of these proteins and their assembled complexes (Ppil4, Smarcal1 and Trip12) as interactors of USP7 TRAF domain. To confirm that these candidate substrates interact with full-length USP7, we used HEK293T cells to express USP7 tagged with streptavidin-binding peptide and FLAG (SFB) prior to pull-down with streptavidin beads (Figure 5A). Immunoblotting detection of endogenous candidate substrates showed that Trip12, Smarcal1, Ubxn7, Ppil4, Ring1 and Fam172a co-precipitated with full-length USP7. Trip12 and Ppil4 displayed the strongest interaction with USP7, as demonstrated by their enrichment in the pull-down fraction compared to in the input lysate.

**Figure 5.**
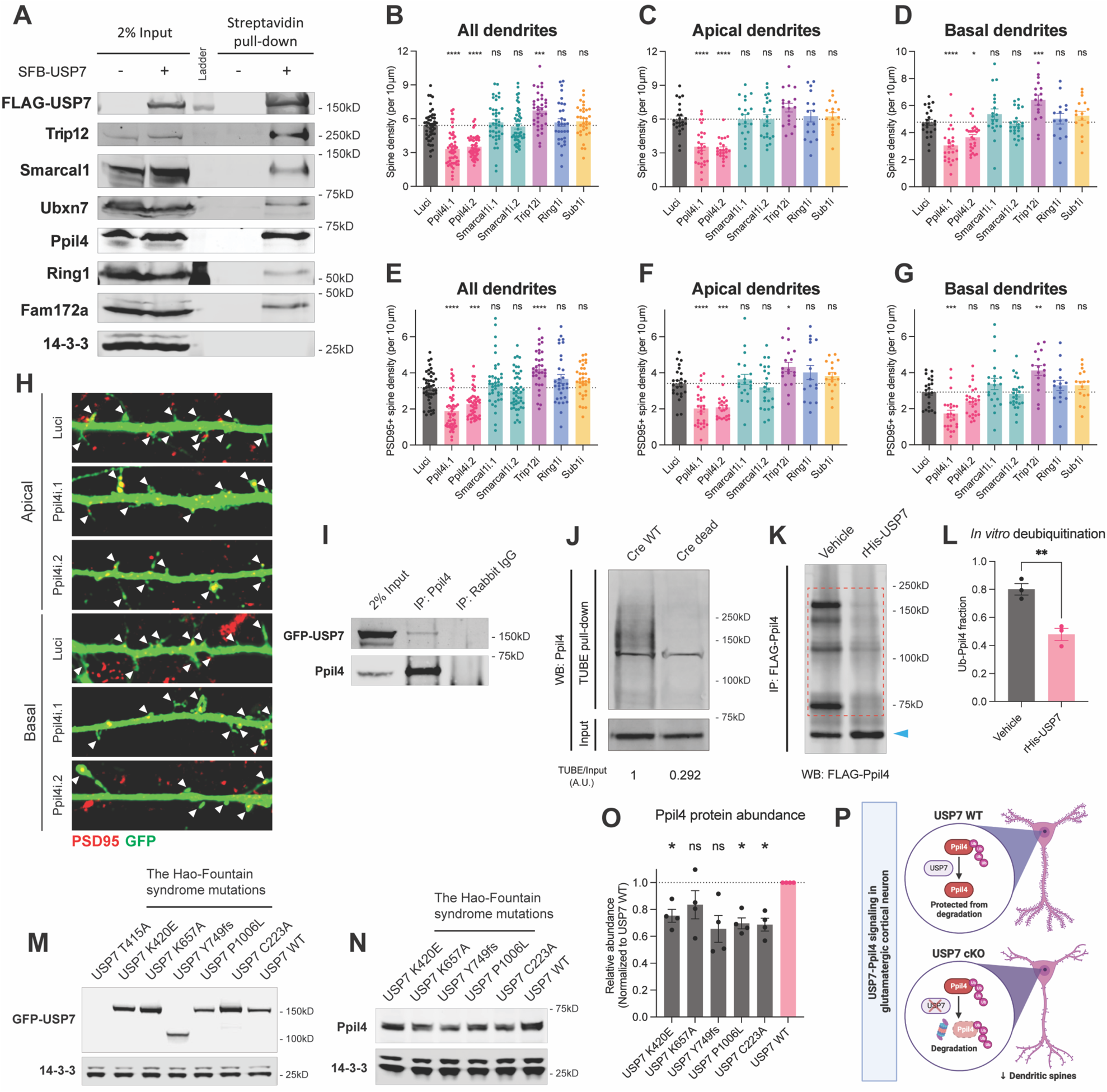
USP7 targets and deubiquitinates Ppil4 to promote synaptogenesis. (A) Streptavidin pull-down of HEK293T lysate with overexpression of Streptavidin-binding peptide-FLAG (SFB)-tagged USP7 followed by western blot of multiple endogenous candidate substrates. 2% input lysate is shown on the left. (B, C, D) Density of dendritic spine on apical, basal and all dendrites of pyramidal-shaped neurons in (H). Two independent lentiviral shRNA knocking down Ppil4 reduced spine density compared with control shRNA targeting Luciferase (Luci). Number of neurons (apical dendrties) = 24, 27, 23, 20, 23, 18, 16 & 17 for Luci, Ppil4i.1, Ppil4i.2, Smarcal1i.1, Smarcal1i.2, Trip12i, Ring1i & Sub1i. Number of neurons (basal dendrties) = 22, 24, 26, 20, 24, 17, 15 & 16 for Luci, Ppil4i.1, Ppil4i.2, Smarcal1i.1, Smarcal1i.2, Trip12i, Ring1i & Sub1i. *p<0.05, **p<0.01, ***p<0.001, ****p<0.0001 by Dunnett’s multiple comparison test (compared to Luci). (E, F, G) Density of PSD95+ dendritic spine on apical, basal and all dendrites of cortical neurons in (H) and Figure S10I. Two independent lentiviral shRNA knocking down Ppil4 reduced PSD95+ spine density compared with control shRNA targeting Luciferase (Luci). Number of neurons are the same as in (B, C, D). *p<0.05, **p<0.01, ***p<0.001, ****p<0.0001 by Dunnett’s multiple comparison test (compared to Luci). (H) Representative images show the effect of Ppil4 knockdown on dendritic spines of cortical neurons at DIV18. PSD95 puncta decorating the dendritic spines were immunostained along with GFP. Arrows show dendritic spines. Scale bar 5µm. (I) Immunoprecipitation of endogenous Ppil4 in HEK293T lysates followed by western blot of GFP-USP7 and Ppil4. 2% Input lysate is shown on the left. GFP-USP7 was co-immunoprecipiated with Ppil4. (J) Tandem Ubiquitin Binding Entities (TUBE) pull-down of neuronal lysate with USP7 loss driven by lentiviral Cre, followed by immunoblotting of Ppil4. Input lysates are shown at the bottom. Endogenous Poly-Ubiquitinated Ppil4 increased with USP7 loss (Cre WT). A.U. = aribitary unit. (K) *In vitro* deubiquitination assay using FLAG-immunoprecipitated Ppil4 from HEK293T cells. Incubation with recombinant His-USP7 at 37°C for 1 hour converted high-molecular weight ubiquitinated Ppil4 (dashed rectangle) into low-molecular weight Ppil4 (arrow). (L) Densitometric quantification of high-molecular weight Ppil4 over all Ppil4 signal in (K) over 3 biological replicates. **p<0.01 by 2-tail unpaired t-test. (M) Western blot of USP7 with mutations from the Hao-Fountain syndrome patients in HEK293T cells. USP7 C223A is catalytically dead. (N) Western blot of endogenous Ppil4 in HEK293T cells expressing different USP7 variants. (O) Densitometric quantification of Ppil4 signal as in (N) over 4 biological replicates. Repeated measurement one way ANOVA. *p<0.05 by Dunnett’s multiple comparison test. (P) Schematic of USP7-Ppil4 signaling pathway regulating dendritic spine density in glutamatergic cortical neuron. Data are presented as mean ± SEM. See also Figure S10.

Reduction in dendritic spine density was a prominent phenotype in neurons with USP7 loss (Figure 3). Therefore, we asked whether select candidate substrates might regulate the morphogenesis of dendritic spines. We tested the effect of knockdown of the candidate substrates at DIV4 using lentiviral shRNA in wild-type primary cortical neurons (validated in Figure S10A-E) before assessing dendritic spine formation at DIV18 (Figure 5B-D). We found that knockdown of Ppil4 using two independent shRNAs significantly decreased the density of dendritic spines (Figure 5B-D and 5H). Further immunofluorescence analysis of PSD95, a major postsynaptic component of dendritic spines, confirmed the reduction of PSD95-positive spines in neurons subjected to Ppil4 knockdown (Figure 5E-H and S10F-H). These observations suggest that Ppil4 is required for the formation of mature, stable synapses. Surprisingly, knockdown of Trip12, a ubiquitin ligase associated with intellectual disability^25,26^, led to an increase in spine density (Figure 5B-H and Figure S10F-I). Knockdown of other candidate USP7 substrates, including the SWI/SNF component Smarcal1 had little to no effect on spine density (Figure 5B-H and Figure S10F-I). Taken together, these results suggest that Ppil4 promotes synaptogenesis in cortical neurons, and depletion of Ppil4 is sufficient to phenocopy the effects of USP7 loss.

We next determined whether Ppil4 is a direct target of USP7-mediated deubiquitination. First, we found that endogenous Ppil4 interacts with GFP-tagged USP7 in HEK293T cells by co-immmunoprecipitation (Figure 5I). Using the Tandem Ubiquitin Binding Entities (TUBE) assay^27^, we purified endogenous polyubiquitinated proteins at DIV5 from lysates of primary cortical neurons of genotype *Usp7*^fl/fl^; *Trp53*^+/−^ transduced with Cre lentivirus at DIV3. Subsequent immunoblotting analyses showed that, compared to catalytically dead Cre, wild-type Cre lentiviral transduction triggered a dramatic increase of high-molecular weight forms of Ppil4, indicative of ubiquitination of Ppil4 upon USP7 loss (Figure 5J). Finally, we found that recombinant USP7 was sufficient to deubiquitinate FLAG-Ppil4 immunopurified from HEK293T cells in an *in vitro* cell-free system (Figure 5K & 5L). Collectively, these observations demonstrate that Ppil4 is a bona fide target of USP7 deubiquitinase activity, and that USP7 and Ppil4 regulate the formation of dendritic spines and synapses of glutamatergic neurons (Schematic as Figure 5P).

Lastly, to examine the relevance of Ppil4 as a USP7 substrate in the context of the Hao-Fountain syndrome, we asked whether reported patient mutations in USP7 disrupt the USP7-Ppil4 signaling pathway. Among the 5 USP7 patient mutations^28–31^ (T415A, K420E, K657A, Y749fs and P1006L) we tested, four were stably expressed in HEK293T cells (Figure 5M). Subsequent immunoblotting analyses showed that the protein abundance of endogenous Ppil4 was reduced in HEK293T cells expressing either patient USP7 mutants or catalytically dead USP7 (C223A), compared to cells expressing wild-type USP7 (Figure 5N & 5O). These results suggest that USP7 mutants in the Hao-Fountain syndrome patients lead to destablization of Ppil4, similar to the catalytically inactive USP7 mutant.

## Discussion

In this study, we have discovered novel roles and underlying mechanisms for the Hao-Fountain syndrome protein USP7 in brain development. Conditional knockout of the deubiquitinase USP7 in post-mitotic glutamatergic neurons of the forebrain leads to deficits in mice, from sensorimotor dysfunction to abnormalities of cognition and social behavior, reminiscent of phenotypes in the Hao-Fountain syndrome. In developmental studies, conditional USP7 loss leads to perinatal apoptosis in post-mitotic glutamatergic neurons, and genetic deletion of p53 rescues the USP7 loss-induced apoptosis. Deletion of p53 also rescues impairments in visual acuity and auditory cue fear conditioning in *Usp7* cKO mice. Strikingly, however, loss of p53 fails to rescue most behavioral deficits or the brain volume reduction upon loss of USP7, indicating that USP7 regulates brain development and function predominantly in a p53-independent manner. Accordingly, USP7 loss reduces the amounts of synaptic proteins in the cerebral cortex and density of dendritic spines in cortical pyramidal neurons in the background of p53 deficiency. In immunoprecipitation followed by mass spectrometry analyses in primary neurons, we uncover the interaction of the USP7 TRAF domain, which binds substrates in proliferating cells, with ribonucleoprotein and RNA spliceosome complex proteins in neurons. Importantly, in quantitative TMT-proteomics analyses, the abundance of the ribonucleoprotein and spliceosome proteins covaries with USP7 in control and USP7-deleted primary cortical neurons. Among the spliceosome proteins, we identify the RNA splicing factor Ppil4 as a novel substrate of USP7 that is deubiquitinated by USP7. Knockdown of Ppil4 phenocopies the effect of USP7 loss on dendritic spine morphogenesis in cortical neurons. Our findings define a novel ubiquitin signaling link between the deubiquitinase USP7 and the splicing factor Ppil4 that operates independently of p53 to drive synapse morphogenesis in the developing brain, with ramifications for our understanding of the Hao-Fountain syndrome.

Our findings have significant implications for studies of ubiquitin signaling networks as well as neurodevelopmental disorders of cognition. The USP7-p53 pathway has been extensively characterized in the context of cancer biology^10,11,32^. Our finding that USP7 loss induces p53-dependent apoptosis in glutamatergic neurons broadens the relevance of the p53 pathway to post-mitotic cells. However, despite the requirement for p53 in cortical neuron apoptosis in *Usp7* cKO mice, the majority of the behavioral deficits in *Usp7* cKO mice persist independently of p53 deletion. Consistently, impairment of dendritic spine morphogenesis upon USP7 loss occurs independently of p53 loss. The loss of p53 also fails to reverse the reduction in brain volume in *Usp7* cKO mice, suggesting that USP7 may regulate brain size in a p53-independent manner. Because USP7 loss has little effect on soma size and dendrites (Figure S6), our data suggest that the reduction of dendritic spines in USP7-deficient neurons may contribute to the reduced volume of the cerebral cortex^33^. Whereas dendritic spines house excitatory, but not inhibitory, synapses^34^, our proteomic analyses in the cerebral cortex reveal that down-regulation of both excitatory and inhibitory synapse proteins occurs upon USP7 depletion. This observation raises the question of whether USP7 also regulates the development of inhibitory synapses in the brain. Because abnormalities of synapse development are thought to underlie the pathogenesis of neurodevelopmental disorders of cognition^35–37^, USP7 regulation of dendritic spine morphogenesis in glutamatergic neurons may play a key role in the pathogenesis of the Hao-Fountain syndrome.

One key finding in this study is the identification of the RNA splicing factor Ppil4 as a novel substrate of USP7 in post-mitotic neurons, which bears broad implications for studies of the major deubiquitinase USP7. First, although much of what has been learned about USP7 has been in the context of cancer biology where USP7 is thought to promote tumorigenesis through the deubiquitination of the ubiquitin ligase Mdm2 and consequent degradation of p53^10,11^, the identification of Ppil4 as a novel substrate elucidates a novel mechanism in post-mitotic neurons in the developing brain with relevance to our understanding of the Hao-Fountain syndrome. Second, by deubiquitinating and consequent regulation of the levels of the protein Ppil4, our findings suggest that USP7 may regulate RNA splicing in neurons, which will be the subject of future studies. Our findings in interaction and quantitative TMT-proteomics analyses revealing that USP7 interacts with and regulates the abundance of small nuclear ribonucleoproteins, the RNA helicase protein Dhx, and the RNA regulatory protein Fam172a^38,39^, further support a role for USP7 in the control of RNA processing. Recent studies suggest that Ppil4 is recruited to active spliceosomes to facilitate catalytic activation of the spliceosome by positioning the key RNA helicase Prp2 downstream of the branching site adenosine of introns^40,41^. It will be important to determine how USP7 regulation of Ppil4 ubiquitination might influence the catalysis of RNA splicing in neurons. Third, the observation that Ppil4 is required for dendritic spine morphogenesis, phenocopying the effect of USP7 loss in cortical neurons, raises the important question of whether the impairment of RNA splicing might contribute to the pathogenesis of the Hao-Fountain syndrome. Because alternative splicing in neurons is frequently associated with the regulation of synaptic proteins including adhesion molecules and scaffold proteins^42–44^, it may be interesting in future studies to test whether the USP7-Ppil4 ubiquitin signaling pathway regulates the splicing of these molecules to influence dendritic spinogenesis. Finally, whereas the discovery of the USP7-Ppil4 ubiquitin signaling link plays a critical role in the brain, our findings also raise the question of whether this mechanism operates in proliferating cells and in the context of cancer.

Besides Ppil4, our mass spectrometry analyses also reveal the interactome of USP7 and proteomic dynamics upon acute USP7 loss for the first time in post-mitotic neurons. To our knowledge, this represents the first comprehensive characterization of potential substrates of a deubiquitinase in neurons. Notably, we observed that multiple ubiquitin ligases interact with USP7, and the protein abundance of these ubiquitin ligases is reduced upon USP7 loss. Ubiquitin ligases characteristically undergo self-ubiquitination and self-degradation^11,45,46^. Therefore, USP7 might function as a positive regulator of these ubiquitin ligases by limiting their self-ubiquitination and degradation. Interestingly, among these ubiquitin ligases, Trip12 and Huwe1 are both associated with intellectual disability^25,26,47,48^, thus suggesting that the regulation of these ubiquitin ligases by USP7 may bear potential relevance to the Hao-Fountain syndrome. Furthermore, our proteomic and biochemical analyses have also identified several chromatin and transcription regulators as novel candidate substrates of USP7 in neurons. In particular, the chromatin remodeling SWI/SNF class enzyme Smarcal1 interacts with USP7 and is stabilized by USP7, suggesting a role for USP7 in regulating chromatin accessibility.

The characterization of the behavioral deficits in *Usp7* cKO mice reveals that these mice display several phenotypes reminiscent of the clinical features of patients with the Hao-Fountain syndrome, including impairment in visual function, motor strength, gait, cognition, and aggression. Although patients with the Hao-Fountain syndrome typically exhibit germline heterozygous USP7 deletion or mutation^8,9^, the phenotypic similarities between patients and our *Usp7* cKO mice suggest a shared biology for USP7 in both humans and mice. Our data suggest that USP7 dysfunction in glutamatergic excitatory neurons might be relevant to the pathogenesis of the Hao-Fountain syndrome. In particular, *Usp7* cKO mice display a tail-biting behavior, not seen to our knowledge in other mouse models, which may bear implications for understanding of aggression in the Hao-Fountain syndrome. Furthermore, our results that both USP7 loss and USP7 patient mutations reduce the level of the substrate Ppil4 suggest that USP7 loss of function may represent a shared mechanism between *Usp7* cKO mice and patients with the Hao-Fountain syndrome. Collectively, our findings suggest that loss of USP7 function in excitatory glutamatergic neurons in the forebrain triggers phenotypes that might be relevant to the Hao-Fountain syndrome. In future studies, it will also be crucial to determine the effect of USP7 deletion in other cell types, including interneurons and glial cells in the brain, and their contributions to the pathogenesis of the Hao-Fountain syndrome.

In conclusion, we have elucidated the roles and mechanisms of the Hao-Fountain syndrome protein USP7 in the mammalian brain. Loss of USP7 in glutamatergic excitatory neurons in the forebrain triggers mouse phenotypes with similarities to phenotypes in the Hao-Fountain syndrome patients. We have found that USP7 regulates the development and function of the brain via p53-dependent and predominantly p53-independent mechanisms, leading to the discovery of the USP7-Ppil4 ubiquitin signaling link that plays a key role in dendritic spine morphogenesis in the developing brain. Beyond advancing our understanding of the major deubiquitinase USP7 in normal brain development and function, our study provides clues for the identification of biomarkers and potential treatment of patients with the Hao-Fountain syndrome.

### Limitations of the study

Although our study provides fundamental new insights into the functions and mechanisms of USP7 in the developing mammalian brain, a number of limitations should be considered. Whereas mouse genetics provides a rigorous approach to defining functions and mechanisms of the Hao-Fountain syndrome protein USP7 in a living organism, the relevance of mice to studies of human disorders remains a limitation. In the future, it will be worthwhile to extend these studies to human model systems including induced pluripotent stem cell (iPSC)-derived cells and tissues. In our mechanistic studies, although we focused on Ppil4 as a key target of USP7, in the future, it will be important to extend our studies to the investigation of additional candidate substrates. Importantly, ubiquitination influences protein degradation as well as other aspects of protein function independently of protein turnover. Our TMT proteomics analyses and follow-up biochemical experiments focused on the degradation pathway. Hence, it remains to be determined whether USP7 impacts signaling pathways in a manner independent of its effects on protein turnover.

## Acknowledgements

The authors would like to thank the Kim lab and Bonni lab members for critical feedback. The authors also thank A. Yen for the help with AAV injection and N. Mosammaparast and N. Tsao for the help with TUBE assay. This work was supported by: NIH/NINDS R01NS051255 (to A.B. and A.H.K), R01 NS111014 (to H.Y.), NIH/NIGMS R01 GM132129 (J.A.P.) and GM67945 (S.P.G.), a McDonnell Center for Systems Neuroscience grant award (to A.H.K.), the Christopher Davidson and Knight Family Fund (to A.H.K.), and the Duesenberg Research Fund (to A.H.K.). This work was also supported by the Washington University Center for Cellular Imaging (S10 OD21629-01A1-WUCCI) and Hope Center Viral Vectors Core at Washington University School of Medicine.

## Author contributions

HC, CJF, AB and AHK designed the project and all the experiments. DM and SPG performed the TMT experiments. JP and SPG performed the IP-MS experiment. AT and CMY performed and analyzed the behavioral experiments. AH analyzed the *in vivo* apoptosis and dendritic spine shape experiment. THL and SKS performed the *ex vivo* MRI and volumetric measurement. HY contributed to DNA cloning. WG provided the USP7^fl/fl^ mouse line. HC performed and analyzed the rest of the experiments and wrote the manuscript. CJF, AB and AHK edited the manuscript.

## STAR Methods

### Key resources table

| REAGENT or RESOURCE | SOURCE | IDENTIFIER |
| --- | --- | --- |
| **Antibodies** |  |  |
| Rabbit anti-USP7 | Bethyl labs | Cat# A300-033A |
| Rabbit anti-USP7 | Bethyl labs | Cat# IHC-00018 |
| Mouse anti-NeuN | Millipore | Cat# MAB377 |
| Rat anti-Ki67 (SolA15) | Thermo Fisher Scientific | Cat# 13-9910-82 |
| Rabbit anti-Cleaved caspase 3 | Cell signaling | Cat# 9661 |
| Chicken anti-GFP | Abcam | Cat# ab13970 |
| Rabbit anti-GFP | Homemade | N/A |
| Mouse anti-mCherry (1C51) | Novus Biological | Cat# NBP1-96752 |
| Mouse anti-Cre (2D8) | Millipore | Cat# MAB3120 |
| Rabbit anti-Cre | Millipore | Cat# 69050 |
| Mouse anti-Myc (9e10) | Santa Cruz Biotechnology | Cat# sc-40 |
| Mouse anti-FLAG (M2) | Sigma | Cat# F3165 |
| Mouse anti-FLAG M2 agarose beads | Sigma | Cat# A2220 |
| Rabbit anti-HA (C29F4) | Cell Signaling | Cat# 3724 |
| Rabbit anti-Map2 | Abcam | Cat# ab32454 |
| Rabbit anti-Phip | Bethyl labs | Cat# A302-054A |
| Rabbit anti-Xiap | Bioss Antibodies | Cat# bs-1281R |
| Rabbit anti-Fto | Abclonal | Cat# A1438 |
| Rabbit anti-Rnf220 | Abclonal | Cat# A15853 |
| Rabbit anti-Cfdp1 | Bethyl labs | Cat# A305-624A |
| Rabbit anti-Phc2 | Abclonal | Cat# A14727 |
| Rabbit anti-Ppil4 | Bethyl labs | Cat# A304-965A |
| Rabbit anti-Ppil4 | Sigma | Cat# HPA031600 |
| Rabbit anti-Gpd1 | Abclonal | Cat# A5715 |
| Rabbit anti-Smarcal1 | Millipore | Cat# ABE1836 |
| Rabbit anti-Ubxn7 | Bethyl labs | Cat# A303-865A |
| Rabbit anti-Rnf2 | Abclonal | Cat# A5563 |
| Rabbit anti-Trip12 | Abclonal | Cat# A9958 |
| Rabbit anti-Fam172a | Novus Biological | Cat# NBP1-86774 |
| Rabbit anti-Sub1 | Cohesion Biosciences | Cat# CQA1533 |
| Rabbit anti-Chfr | Abclonal | Cat# A10447 |
| Mouse anti-Ruvbl2 | Santa Cruz | Cat# sc-374135 |
| Mouse anti-Ring1 (8C12F4) | Novus Biological | Cat# NBP2-37370 |
| Mouse anti-Gfap | Millipore | Cat# MAB3402 |
| Mouse anti-14-3-3 | Santa Cruz Biotechnology | Cat# sc-25276 |
| Mouse anti-Lamin A/C (4C11) | Cell signaling | Cat# 4777 |
| Mouse anti-PSD95 (K28/43) | NeuroMab | Cat# 75-028 |
| Normal Rabbit IgG | Cell signaling | Cat# 2729 |
| **Bacterial and virus strains** |  |  |
| Lentivirus | This study | N/A |
| AAV PHP.eB | Hope Center Viral Vectors | N/A |
|  | Core at Washington University |  |
| **Chemicals, peptides, and recombinant proteins** |  |  |
| MG132 | Sigma | Cat# 474790 |
| Bradford reagent | Bio-Rad | Cat# 5000006 |
| Protease inhibitor cocktail | Sigma | Cat# P8340 |
| Phosphatase inhibitor mini-tablet | Thermo Fisher Scientific | Cat# A32957 |
| Hoechst 33258 solution | Sigma | Cat# 94403 |
| 3x FLAG peptide | APExBio | Cat# A6001 |
| Dynabeads™ MyOne™ Streptavidin T1 | Invitrogen | Cat# 65601 |
| TUBE 2 magnetic beads | LifeSensors | Cat# UM-0402M |
| Recombinant His-USP7 | R&D Systems | Cat# E-519 |
| **Deposited data** |  |  |
| IP mass spectrometry data | ProteomeXchange | PXD043225 |
| TMT proteomics data | ProteomeXchange | PXD043848 |
| TMT proteomics data | ProteomeXchange | PXD043849 |
| **Experimental models: cell lines** |  |  |
| HEK293T/HEK293LE | Clontech | Cat# 632180 |
| **Experimental models: organisms/strains** |  |  |
| NEX-cre mice on C57BL/6J | Goebbels et al., 2006 | N/A |
| USP7 fl/fl mice on hybrid C57BL/6J and 129Sv | Kon et al., 2011 | N/A |
| USP7 fl/fl mice on C57BL/6J | This study | N/A |
| p53 KO mice on C57BL/6J | The Jackson Laboratory | Cat# 002101 |
| Wild-type outbred mice on CD1 | Charles River | Cat# 022 |
| Wild-type inbred mice on C57BL/6J | The Jackson Laboratory | Cat# 000664 |
| Wild-type outbred Long-Evans rats | Charles River | Cat# 006 |
| **Oligonucleotides** |  |  |
| See Table S2 | N/A | N/A |
| **Recombinant DNA** |  |  |
| pCMV-myc3-Mdm2 | Addgene | Cat# 20935 |
| pAAV-hSyn1-EGFP-P2A-EGFPf-WPRE-HGHpA | Addgene | Cat# 74513 |
| pMH-SFB-USP7 | Addgene | Cat# 99393 |
| pEGFP-USP7 | Anckar et al., 2015 | N/A |
| pEGFP-USP7 1-209 | This study | N/A |
| pEGFP-USP7 55-209 (TRAF) | This study | N/A |
| pEGFP-USP7 C223A (catalytically dead) | This study | N/A |
| pEGFP-USP7 T415A | This study | N/A |
| pEGFP-USP7 K420E | This study | N/A |
| pEGFP-USP7 P657A | This study | N/A |
| pEGFP-USP7 Y749fs | This study | N/A |
| pEGFP-USP7 P1006L | This study | N/A |
| pHAGE-hSyn-FLAG-USP7 55-209 (TRAF) | This study | N/A |
| FCIV.1-Cre-IRES-mCherry | This study | N/A |
| FCIV.1-Cre Y331F-IRES-mCherry (Cre dead) | This study | N/A |
| pcDNA3.1-FLAG-Ppil4 | This study | N/A |
| **Software and algorithm** |  |  |
| ImageJ-Fiji | National Institutes of Health, USA | https://github.com/fiji/fiji |
| Zen | Zeiss | N/A |
| Imaris | Oxford Instruments | N/A |
| Image Studio | Li-Cor | N/A |
| Proteomics software pipeline #1 | SEQUEST | http://fields.scripps.edu/yates/wp/?page_id=17 |
| Proteomics software pipeline #2 | Comet | https://uwpr.github.io/Comet/ |
| GSEA (v4.3.2) | UC San Diego, Broad Institute | https://www.gsea-msigdb.org/gsea/index.jsp |
| Cytoscape (v3.9.1) | National Institute of General Medical Sciences | https://cytoscape.org/ |
| ITK-SNAP (v3.8) | NYU, Tanden School of Engineering | www.itksnap.org |
| Prism (v.9) | GraphPad | N/A |
| SPSS (v.26) | IBM | N/A |
| R | The R project | N/A |

### Data and materials availability

All mouse lines, cell lines, purified protein or any other reagent generated or used in this study will be made available to researchers who are interested. The proteomics mass spec datasets (IP-MS, *in vivo* TMT and *in vitro* TMT) were deposited at ProteomeXchange via the PRIDE database and will be publicly available upon publication. Accession numbers for these datasets are listed in the Key resources table. Any additional information required to reanalyze the data reported in this paper is available from the corresponding authors upon request.

### Experimental model and subject details

#### Mice

Animals were cared for in accordance with NIH guidelines. All experimental methods were approved by Washington University Institutional Committee on the Use and Care of Animals under the protocol number 20180303 and 210083. Animals were housed in a 12:12 light:dark cycle. We employed trio breeding and only used unaffected animals as parents. Pups were weaned from P21 to P28 and genotyped by phenol/chloroform DNA extraction of tail or toe (< P7 pups) biopsy followed by PCR. Weaned mice were separated by sex and relocated for breeding from 6 to 24 weeks’ old. Animals were allocated to experimental versus control groups based on genotype. Sex matched-littermate controls were used in all experiments, and the age at which animals were used is reported in figure legends.

The *Usp7* floxed mice were first obtained on C57BL/6J and 129Sv mixed background^15^ and then backcrossed to C57BL/6J background with speed congenics (last generation SNP typing > 99% determined by Dartmouse). The *NEX-cre* mice on C57BL/6J background were obtained from Dr. Hui-Chen Lu’s lab in Indiana University Bloomington, under the permission of Dr. Sandra Goebbels^19^. Only the videotaped *Usp7* cKO mice were generated from the *Usp7* floxed mice on C57BL/6J and 129Sv mixed background while the mice in all experiments were generated from C57BL/6J mice and maintained on C57BL/6J background. *Usp7* cKO mice were separated and single-caged once they were found to bite their cage mates.

The *Trp53* mutant mice on C57BL/6J background were purchased from the Jackson Laboratory (Cat#002101). Consistent with the description from the Jackson Laboratory, female *Trp53* ^−/−^ mice were born under the Mendelian ratio and exhibited young adult lethality. Therefore, only male mice with *Trp53* ^−/−^ were used for breeding and *in vivo* experiments. For cortical neuron culture, *Usp7* ^fl/fl^; *Trp53* ^−/−^ male mice crossed with either *Usp7* ^fl/fl^; *Trp53* ^−/−^ or *Usp7* ^fl/fl^; *Trp53* ^+/−^ female mice to produce embryos with *Usp7* ^fl/fl^; *Trp53* ^−/−^ or *Usp7* ^fl/fl^; *Trp53* ^+/−^. As p53 deficiency increases the propensity to grow tumor, mice and embryos with observed tumor bumps were checked and excluded from all experiments.

The wild-type CD1 mice with timed E16 pregnancy for the *in vitro* apoptosis analysis, and the Long-Evans rat with timed E17 pregnancy for the IP-MS analysis were both purchased from Charles River. No breeding were made with these animals.

#### HEK293T cell line

Human embryonic kidney 293T (HEK293T/HEK293LE; Clontech 632180) cells were cultured in DMEM with 10% fetal bovine serum (Fisher #10082-147) and penicillin/streptomycin (Life Technologies #15140122). For lentiviral packaging, DNA/PEI (Polysciences) complexes were suspended in Opti-MEM medium (Gibco) and transfected into HEK293T cells with about 80% confluency. The transfer plasmid was mixed with packaging plasmid psPAX2 and envelope plasmid pCMV-VSVG. The medium was replaced on the next day and the new medium containing lentivirus was collected after 3 days. Cell debris was filtered from the collected medium and Lenti-X Concentrator (Clontech) was then used to precipitate the lentivirus in the medium. After incubation at 4°C overnight to 3 days, virus-containing pellets were collected by centrifuging at 1500g for 45 min at 4°C, resuspended in phosphate-buffered saline (PBS), and stored at −80°C in aliquots.

#### Cortical neuron culture

For IP-MS and immunocytochemistry of USP7 TRAF domain, cortical neurons were cultured from E17 Long-Evans rat embryos. For *in vitro* apoptosis analysis, cortical neurons were cultured wild-type E16 CD1 mouse embryos. For dendrite morphology analysis and TUBE assay, cortical neurons were cultured from E15 *Usp7* ^fl/fl^; *Trp53* ^+/−^ embryos on C57BL/6J background. For *in vitro* TMT proteomics, cortical neurons were cultured from E15 *Usp7* ^fl/fl^; *Trp53* ^−/−^ embryos on C57BL/6J background. For spine analysis with RNAi knockdown, cortical neurons were cultured from E15 C57BL/6J wild-type embryos.

To prepare cortical neurons (previously described as Spiegel et al. 2014^49^), embryonic cortices were dissected in 1X HBSS and digested with 2.5% Trypsin (Gibco) at 37°C. DNase (Roche) was added to digest free-floating DNA from dead cells. Trypsin-digested tissue was further triturated with pipette acid to fully dissociate cells. Dissociated neurons were seeded onto tissue culture dishes pre-coated with poly-L-lysine (0.1mg/mL) at a density of 1.2×10^5^ cells/cm^2^. Neurons were first cultured with neurobasal medium with 1% fetal bovine serum (2% for rat neuron), 0.5mM glutamine, B27 Supplement (Gibco) and penicillin/streptomycin (Life Technologies #15140122) at 37°C with 5% CO^2^. On DIV4, DIV8 and DIV12, neurons were fed with half fresh medium without serum. Every day after DIV14, one fifth of the old medium was replaced with fresh medium without serum to maintain stable medium composition. Neuron viability was confirmed at DIV11 with MTS assay.

### Method details

#### General design of behavioral tests

All behavior tests were conducted in the Washington University Animal Behavior Core and are approved by the Washington University Institutional Animal Care and Use Committee. Sex and litter-matched mice with and without p53 heterozygosity were tested separately due to limit of cohort size for each batch of phenotypical analysis. After the event that one genotype of mice were all dead or severely wounded, all mice in the litter were removed from the analysis to reassure all animals are litter-matched. All tests were performed during the light phase of the light/dark cycle by a female experimenter blind to the genotypes. Mice were acclimated to the test room for 30 min prior to testing.

#### Sensorimotor battery

Ledge, platform, pole, and inclined and inverted screen tests were performed to assess sensorimotor function. Time in each task was manually recorded. The average for two trials was used for analyses. Test duration was 60s, except for the pole test, which was extended to 120s. For walking initiation, time for an animal to leave a 21×21cm square on a flat surface was recorded. For ledge and platform tests, the time the animal was able to balance on an acrylic ledge (0.75cm wide and 30cm high), and on a wooden platform (1.0cm thick, 3.0cm in diameter and elevated 47cm) was recorded, respectively. The pole test was used to evaluate fine motor coordination by quantifying time to turn 180° and climb down a vertical pole. The screen tests assessed a combination of coordination and strength by quantifying time to climb up or hang onto a mesh wire grid measuring 16 squares per 10cm, elevated 47cm and inclined (60° or 90°) or inverted.

#### Catwalk

Natural gait parameters were measured using the CatWalk XT gait analysis system (Noldus, Inc). Mice were habituated to the apparatus and testing room the day before testing by placing each mouse on the glass walkway for 5 min to help facilitate movement during the test session. Testing was conducted with the room lights off. The following day, mice were placed on the 8-cm wide glass walkway and allowed to walk naturally until 3 recordings were collected meeting compliance criteria of less than 60% speed variability. A camera below the glass surface recorded the paw placement and intensity of each paw as the mouse moved across the walkway. Run speed, swing speed, stride length, and mean intensity of paw placement were analyzed to assess differences in gait.

#### Virtual Optomotor System (VOS)

Visual acuity was assessed by recording Spatial frequency thresholds, in cycles per degree (c/d) using the mouse OptoMotry HD System (Cerebral Mechanics Inc.). This is a virtual-reality system designed to quantify visuomotor behavior^50^. Mice were allowed to acclimate to procedure room 30 minutes prior to testing. Each mouse was placed onto a platform in the center of the VOS chamber, and the experimenter tracked the movement of the animal’s head during the course of the test. VOS chamber consists of four computer screens surrounding the platform with a camera to capture the OMR from above. Drift speed was set at 12.5, contrast 100 degree. The OptoMotry software uses a staircase method to determine at which threshold the optomotor reflex (OMR) is no longer detected by the experimenter. Experimenter remained blind to the direction of virtual movement.

#### Spontaneous alternation Y-maze

Testing was conducted according to our previously published procedures^51^. Briefly, this involved placing a mouse in the center of a Y-maze that contained three arms that were 10.5 cm wide, 40 cm long and 20.5 cm deep where an arm was oriented at 120° with respect to each successive other arm. Mice were allowed to explore the maze for 8 min and entry into an arm was scored only when the hindlimbs had completely entered the arm. An alternation was defined as any three consecutive choices of three different arms without re-exploration of a previously visited arm. The number of alternations and arm entries along with the percentage of alternations, which was determined by dividing the total number of alternations by the total number of entries minus 2, then multiplying by 100 was analyzed as a measure of spontaneous alternation.

#### Acoustic startle response/auditory function

Acoustic startle responses were measured using Kinder Scientific Startle Reflex equipment and software (Kinder Scientific, Inc) Auditory function was assessed by measuring startle response to auditory stimuli pulses (40 ms broadband burst) presented at 80dB, 90dB, 100dB, 110dB, and 120dB above background noise (65dB). Each trial type was presented 10 times in a random order over a 20 min session, with inter-trial-intervals ranging from 5-15 sec between trials. Beginning at stimulus onset, 1 ms force readings were obtained to record an animal’s startle amplitude. The average force response to each sound pressure level was analyzed as a measure of auditory function.

#### Conditioned fear

Two distinct chambers were used to train and test mice (26×18×18 cm high) (Med-Associates, St. Albans, VT) which were easily distinguished by different olfactory, visual, and tactile cues present in each chamber. On day 1, each mouse was placed into the conditioning chamber (peppermint odor) for 5 min and freezing behavior was quantified during a 2 min baseline period. Freezing (no movement except that associated with respiration) was quantified using FreezeFrame image analysis software (Actimetrics, Evanston, IL) which allows for simultaneous visualization of behavior while adjusting for a “freezing threshold” during 0.75 s intervals. After baseline measurements, a conditioned stimulus (CS) consisting of an 80 dB tone (white noise) was presented for 20 sec followed by an unconditioned stimulus (US) consisting of a 1 s, 1.0 mA continuous foot shock. This tone-shock (T/S) pairing was repeated each minute over the next 2 min, and freezing was quantified after each of the three tone-shock pairings. Twenty-four hours after training, each mouse was placed back into the original conditioning chamber to test for fear conditioning to the contextual cues in the chamber. This involved quantifying freezing over an 8 min period without the tone or shock being present. Twenty-four hours later, the mice were evaluated on the auditory cue component of the conditioned fear procedure, which included placing each mouse into the other chamber containing distinctly different cues (coconut odor). Freezing was quantified during a 2 min “altered context” baseline period as well as over a subsequent 8 min period during which the auditory cue (CS) was presented. Shock sensitivity was evaluated following completion of the conditioned fear test.

#### Biting behavior

To quantitate biting behavior, Usp7 cKO or Usp7 cKO; Tp53 +/− mice were weaned after 3 weeks old and co-housed with their littermates. Male and female mice were separated and housed in a regular 12:12 light:dark cycle. Tail wounds were checked every week until 12 weeks old. Once tail wounds were found, the aggressors were marked and separated to avoid more biting. Victim mice with severe tail wound were euthanized and photographed. To video record the biting behavior, P33 aggressor and littermate were placed onto an ice cream box lid for free exploration.

#### Calcium phosphate sparse transfection

Primary cortical neurons were transfected as previously described^52^. Briefly, transfection mix was prepared by mixing 50ng/μL plasmid DNA, 0.25mM CaCl_2_ and Hepes Buffered Saline (Invitrogen) and the DNA-calcium phosphate precipitate was made by pipetting up and down 10 times. Neurons were washed 3 times and starved with fresh serum-free medium at 37°C for 30min, and then 40μL of the transfection mix was dropped into each well in a 24-well plate (80μL for 12-well plate) and incubate at 37°C for 20-30 min. To remove DNA-calcium phosphate precipitate and avoid toxicity, neurons were then washed with fresh serum-free medium 3 times again and put back conditioned medium.

#### Immunofluorescence and immunocytochemistry

For immunofluorescence, mice were anesthetized with isoflurane, perfused with 40 m/mL heparin in PBS and 4% paraformaldehyde (> P5) or decapitated (< P5) before brain dissection and immersion fixation of the whole brain in 4% paraformaldehyde overnight at 4°C. Fixed brains were then placed in 30% sucrose in PBS for cryopreservation. After sinking in the sucrose solution, the brains were embedded with mixture of O.C.T. Compound (Sakura Finetek) and 30% sucrose solution (3:7), flash-frozen using liquid nitrogen. With a Leica cryostat, the frozen brains were sectioned at 16μm thickness and placed onto positively charged slides. For spine analysis, the brains were sectioned at 100μm thickness and placed into FD section storage solution (FD NeuroTechnologies). For immunocytochemistry, cells were initially seeded onto HNO_3_-treated coverslips in 12 or 24 well-plates, washed with warm PBS and fixed in 4% paraformaldehyde for 30min on ice. For staining, brain sections or fixed cells were washed with PBS, permeabilized with 0.4% Triton X-100 and blocked with 3% Bovine serum alumin (BSA) and 10% goat serum. The permeabilized and blocked sections or cells was incubated with primary antibody overnight at 4°C, followed by incubation with fluorescence-conjugated secondary antibodies (Invitrogen) for 2 hours at room temperature and Hoechst diluted in PBS at 1:2000. The stained sections or cells were mounted onto slides with Fluoromount-G (SouthernBiotech) or Prolong^TM^ Glass Antifade Mountant (Invitrogen) for spine imaging. Antigen retrieval with citrate acid (0.1 M, pH 6.8 with 0.1% Tween-20) for 15min boiling prior to blocking was required for labeling with anti-Ki67 antibody.

#### *In vivo* apoptosis analysis

Cryosections of P0 mouse brains were immunolabeled with cleaved-caspase 3 (c-cas 3) antibody (Cell Signaling #9961, 1:4000) for apoptosis analysis. Image acquisition and quantitation of the c-cas 3 intensity were blind to mouse genotypes at all steps below. 6-8 consecutive coronal sections per mouse, 3-5 mice per genotype were imaged with Leica DMi8 scanning microscope with instant computational clearing (THUNDER). The imaged sections were selected based on matched hippocampal morphologies across all mice. Segmentation of the cortex to analyze was performed based on Hoechst nuclei counterstain. C-cas 3 integral intensity within the segmented region of each section was measured with ImageJ and normalized to area.

#### *In vitro* apoptosis analysis

Primary cortical neurons from E16 wild-type CD1 mouse embryos were sparsely transfected at DIV3 with U6 empty vector, or U6-driven plasmids of shRNAs targeting USP7 or irrelevant USP30 along with EGFP plasmids. The transfected neurons were fixed at DIV5, 6 and 7, immunolabeled with anti-c-cas 3 (Cell Signaling #9961) and anti-GFP antibody (Abcam #ab13970) and imaged with Zeiss Axio Imager Z2 with ApoTome 2. Image acquisition and quantitation were blind to groups at all steps. Apoptotic cells are counted as GFP+ cells with c-cas3 signal in the cell body and the intensity of apoptosis was quantitated by the percentage of apoptotic cells over all GFP+ cells. The analysis was repeated with three batches of neuron culture and 2-3 wells in 12-well plates were used per group for each culture.

#### Western blot

Cells or cortical tissues were lysed in RIPA buffer (Cell Signaling #9806) or TNE buffer (50mM Tris pH 7.4, 100mM NaCl, 0.1mM EDTA, 1% NP40) mixed with protease inhibitor cocktail (Sigma #P8340). Total protein was quantified with Bradford reagent (Bio-Rad). Lysate samples were separated by SDS-PAGE or LDS-NuPAGE and transferred to 0.2μm nitrocellulose membrane (GE Healthcare) via tank immersion. Membranes were blocked in 5% non-fat Milk in Tris-buffered saline with 0.05% Tween-20 (TBST) at 4°C overnight, incubated with primary antibodies at 4°C overnight, and subsequently with infrared fluorescent secondary antibodies (LI-COR) for 2h at room temperature. Washes with TBST three times were applied to membranes after every antibody incubation step. LI-COR Odyssey CLx imager and LI-COR Image Studio software were used to image the blot at 600 dpi and to determine fluorescence densitometry to quantitate relative protein abundance. The relative abundance of each protein was normalized to loading control proteins (14-3-3 or Lamin A/C) in the same lane. Exposure time was adjusted to avoid pixel saturation.

#### Immunoprecipitation

For immunoprecipitation of GFP-TRAF domain, we generated total cell lysates from HEK293T cells with TNE lysis buffer as described above. 5μM proteasome inhibitor MG132 was applied to enhance the expression of Myc3-Mdm2 in HEK293T cells. 1.5μg rabbit anti-GFP antibody (homemade) was added to 200μL lysate at 2μg/μL total protein concentration determined by Bradford reagent followed by rotation at 4°C overnight. After antibody binding, 18μL of PBS-washed protein G agarose beads (GE Healthcare) were added per immunoprecipitation. Bead binding proceeded for 1 hour at 4°C before 3 times wash with TNE lysis buffer and 3 times wash with cold PBS. 30μL LDS sample buffer was added to beads per immunoprecipitation and heated at 70°C for elution. The precipitated proteins were proceeded to western blot analysis.

For immunoprecipitation of endogenous Ppil4, we generated total cell lysates from HEK293T cells with TNE lysis buffer. 4μg Rabbit anti-Ppil4 antibody (Bethyl) or Rabbit normal IgG control was added to 200μL lysate at 4μg/μL total protein concentration determined by Bradford reagent followed by rotation at 4°C overnight. After antibody binding, 16μL of PBS-washed protein A agarose beads (GE Healthcare) were added per immunoprecipitation. Bead binding proceeded for 1 hour at 4°C before 3 times wash with TNE lysis buffer and 3 times wash with cold PBS. 30μL LDS sample buffer was added to beads per immunoprecipitation and heated at 70°C for elution. The precipitated proteins were proceeded to western blot analysis.

For immunoprecipitation of FLAG-TRAF domain, we generated total neuron lysates from primary rat neurons with the TNE lysis buffer. pHAGE-hSyn-TRAF WT/RDW were transduced into neurons at DIV7 and protein lysates were harvested at DIV10. 20μL PBS-washed M2 anti-FLAG agarose beads (Sigma) were added to 2.5mL lysate at 2μg/μL total protein concentration for 3 hours incubation at 4°C. The beads were washed 3 times with TNE lysis buffer, 3 times with cold PBS, followed by 3 times wash with 200ng/μL 3x FLAG peptides (APExBio) for elution. 1.5% eluted samples were used for LDS-NuPAGE followed by silver stain (Invitrogen) while the rest was proceeded to tricarboxylic acid (TCA) precipitation and LC-MS/MS analysis. 3 biological replicates of neuron plates with TRAF-WT and RDW mutant were used.

#### LC-MS/MS identification of interactors of the USP7 TRAF domain

We resuspended the sample in 20 µL of 8 M urea, 100 mM EPPS pH 8.5. We added 5mM TCEP and incubated the mixture for 15 min at room temperature. We then added 10 mM of iodoacetamide for 15min at room temperature in the dark. We added 15 mM DTT to consume any unreacted iodoacetamide. We added 180µl of 100 mM EPPS pH 8.5. to reduce the urea concentration to <1 M, 1 µg of trypsin, and incubated at 37 C for 6 hrs. The solution was acidified with 2% formic acid and the digested peptides were desalted via StageTip, dried via vacuum centrifugation, and reconstituted in 5% acetonitrile, 5% formic acid for LC-MS/MS processing.

Mass spectrometry data were collected using a Q Exactive mass spectrometer (Thermo Fisher Scientific, San Jose, CA) coupled with a Famos Autosampler (LC Packings) and an Accela600 liquid chromatography (LC) pump (Thermo Fisher Scientific). Peptides were separated on a 100 μm inner diameter microcapillary column packed with ∼20 cm of Accucore C18 resin (2.6 μm, 150 Å, Thermo Fisher Scientific). For each analysis, we loaded ∼1 μg onto the column. Peptides were separated using a 75 min method from 2 to 25% acetonitrile in 0.125% formic acid with a flow rate of ∼300 nL/min. The scan sequence began with an Orbitrap MS^1^ spectrum with the following parameters: resolution 70,000, scan range 300−1500 Th, automatic gain control (AGC) target 1 × 10^5^, maximum injection time 250 ms, and centroid spectrum data type. We selected the top twenty precursors for MS^2^ analysis which consisted of HCD high-energy collision dissociation with the following parameters: resolution 17,500, AGC 1 × 10^5^, maximum injection time 90 ms, isolation window 1.6 Th, normalized collision energy (NCE) 27%, and centroid spectrum data type. The underfill ratio was set at 9%, which corresponds to a 1.5 × 10^5^ intensity threshold. In addition, unassigned and singly charged species were excluded from MS^2^ analysis and dynamic exclusion was set to automatic.

Mass spectra were processed using a SEQUEST-based in-house software pipeline. MS spectra were converted to mzXML using a modified version of ReAdW.exe. Database searching included all entries from the *rat* uniprot database which was concatenated with a reverse database composed of all protein sequences in reversed order. Searches were performed using a 50 ppm precursor ion tolerance. Product ion tolerance was set to 0.03 Th. Carbamidomethylation of cysteine residues (+57.0215Da) were set as static modifications, while oxidation of methionine residues (+15.9949 Da) was set as a variable modification. Peptide spectral matches (PSMs) were altered to a 1% FDR^53,54^. PSM filtering was performed using a linear discriminant analysis, as described previously^55^, while considering the following parameters: XCorr, ΔCn, missed cleavages, peptide length, charge state, and precursor mass accuracy. Peptide-spectral matches were identified, quantified, and collapsed to a 1% FDR and then further collapsed to a final protein-level FDR of 1%. Furthermore, protein assembly was guided by principles of parsimony to produce the smallest set of proteins necessary to account for all observed peptides.

#### TMT mass spectrometry

##### Sample preparation

Preparation of samples prior to mass spectrometry was performed as previously described^56^. In brief, 5 murine cortices were harvested and flash-frozen at P17 and then lysed by bead-beating in lysis buffer (200mM EPPS, pH 8.5 and 8 M urea with protease and phosphatase inhibitors) at maximum speed for four cycles of 30 s each, with 1 min pauses on ice between cycles. For 16-plex experiment, primary mouse cortical neurons were dissociated from E15 USP7 fl/fl; Tp53 −/− embryos on C57BL/6J background. USP7 depletion was induced by lentiviral transduction of wild-type Cre recombinase or catalytically dead Cre as control at DIV3. Four biological replicates of neurons transduced with wild-type Cre and catalytically dead Cre were harvested at DIV6 and DIV8, respectively. For cultured neurons the lysate was passed 10 times through a 21-gauge (1.25 inches long) needle. An equal quantity of protein (100 µg) for each condition (10 cortex slices, 16 samples of transduced neurons) was reduced with 5 mM tris (2-carboxyethyl) phosphine (room temperature, 15 min) before proteins were alkylated with 10 mM iodoacetamide (room temperature, 30 min in the dark). Finally, excess iodoacetamide was quenched with 10 mM dithiothreitol (room temperature, 15 min). Alkylated proteins were isolated by methanol-chloroform precipitation prior to protease digestion. The samples were diluted in lysis buffer to a volume of 100 µL before adding methanol (400 µL), chloroform (100 µL) and water (300 µL) stepwise with vortexing after addition of each solvent. The samples were centrifuged for 1 min at 14,000 g and the aqueous and organic phases were removed, leaving only the protein disk, which was washed with methanol (500 µL) and centrifuged at 21,000 g for 2 min at room temperature. After removal of methanol supernatant, the samples were resuspended in 200 mM EPPS (pH 8.5) containing Lys-C protease (Wako) at a 100:1 protein-to-protease ratio then incubated overnight at room temperature while shaking. Trypsin was then added at a 100:1 protein-to-protease ratio, and the reaction was incubated 6 hr at 37°C. Acetonitrile (30 µL) was added to the digested peptides and vortexed before adding 200 µg TMT reagent (TMT10 or TMTpro16) for a 2:1 TMT-to-peptide ratio. The samples were incubated at room temperature for 1 h and briefly vortexed every 10 min. Once labeling efficiency was confirmed, the reaction was quenched with hydroxylamine to a final concentration of 0.3% (v/v) for 15 min. The TMT-labeled samples were pooled, vacuum-centrifuged long enough to evaporate the acetonitrile, then desalted with a C18 solid-phase extraction (SPE) (Sep-Pak, Waters) and vacuum centrifuged to dryness.

##### Basic reversed-phase fractionation

For basic reversed-phase fractionation, desalted peptides were resuspended in Buffer A (10 mM ammonium bicarbonate, 5% ACN, pH 8), then subjected to a linear gradient from 13% to 42% of Buffer B (10 mM ammonium bicarbonate, 90% ACN, pH 8) on an Agilent 300Extend C18 column (3.5 μm particles, 4.6 mm ID and 250 mm in length). The peptide mixture was fractionated into a total of 96 fractions which were consolidated into 24 super fractions, of which, 12 non-adjacent samples were analyzed by the mass spectrometer. These 12 fractions were vacuum centrifuged until dry, then desalted via StageTip for LC-MS analysis.

##### LC-MS analysis

The LC-MS analysis and protein quantification was slightly different for proteins extracted from the mouse cortex slices versus those from the cultured neurons, reflecting changes in instrumentation over the course of these experiments. Eluted peptides were introduced to an Orbitrap Fusion with FAIMS pro (mouse cortex slices) or Orbitrap Fusion Lumos (cultured neurons) mass spectrometer coupled to a Proxeon EASY-nLC 1200 LC pump (both from ThermoFisher Scientific). Peptides were separated over a 90 min (cortex) or 120 min (neurons) gradient on a 35-40 cm long 100 µm inner diameter column packed with Accucore (2.6 μm, 150Å) resin (ThermoFisher Scientific). For cortex slices the FAIMS voltages were set at −40, −60, and −80V using a top speed method with 1s per CV. Full MS spectra were acquired in the Orbitrap at a resolution of 120,000 and the 10 most intense MS1 ions were selected for MS2 analysis in the ion trap. The isolation width was set to 0.7 Da and isolated precursors were fragmented by CID at a normalized collision energy (NCE) of 35%, and a max injection time of 35 ms (cortex) or 50 ms (neurons). Following acquisition of each MS2 spectrum, for the murine cortex experiment, a synchronous precursor selection (SPS) MS3 scan was collected on the top 10 most intense ions in the MS2 spectrum. For the experiment from cultured neurons, a real-time search (RTS-MS3) strategy was used, where only those SPS ions (up to 10) matching a theoretical peptide spectrum were isolated for MS3 analysis. These SPS-MS3 precursors were fragmented by higher energy collision-induced dissociation (HCD) at an NCE of 65% (TMT10) or 50% (TMTpro) and analyzed using the Orbitrap at 50,000 resolution, with a max injection time of 100 ms (cortex) or 200 ms (neurons).

##### Database searching

Mass spectra were processed using SEQUEST (cortex) or Comet (neurons). Database searching included all entries from the murine Uniprot database downloaded 07/2014 (cortex) or 07/2020 (neurons). Searches were performed using a 50-ppm precursor ion tolerance for total protein-level analysis. The product ion tolerance was set to 0.9 Da (SEQUEST) or 1.005 (Comet). TMT tags (+229.163 Da) or TMTpro tags (+304.2071) on lysine residues and peptide N termini and carbamidomethylation of cysteine residues (+57.021 Da) were set as static modifications, while oxidation of methionine residues (+15.995 Da) was set as a variable modification. Peptide-spectrum matches (PSMs) were identified, quantified, and filtered to a 1% peptide false discovery rate (FDR) and then collapsed further to a final protein-level FDR of 1%. The total summed signal-to-noise (S:N) measurements of the peptides in each condition were used to calculate column normalization factors to account for small differences in sample abundance. Then, for each protein, signal-to-noise (S:N) measurements of the peptides were summed and then normalized to 100 across the all samples to yield a relative abundance measurement.

#### RNA extraction and RT-qPCR

Total RNA from mouse cortical tissues and primary cortical neurons were isolated using QIAshredder and RNeasy Plus Mini Kit (QIAGEN) according to the manufacturer’s instructions. Cortical tissues were homogenized with mortar and pestle on dry ice during lysis. The extracted RNA was reversely transcribed into cDNA by a High-Capacity cDNA Reverse Transcription Kit with RNase Inhibitor (Applied Biosystems). Mixed cDNA, designed primers and Power SYBR Green PCR Master Mix (Applied Biosystems) were loaded to qPCR on a CFX Connect Real-Time PCR Detection System (Bio-Rad) with the program below: hot start, 95°C for 10 min; amplification 40 cycles including denaturation (95°C for 15s), annealing (57°C for 30s), and extension (72°C for 30s). A melting curve analysis (65°C for 5s and then 5s each at 0.5°C increments between 65° and 95°C) was conducted to ensure single amplicon. Glyceraldehyde-3-phosphate dehydrogenase (GAPDH) was used as the internal reference gene and the conversion from cycle number to relative RNA level is performed with Excel 2019 (Microsoft).

#### Streptavidin pull-down

For assessment of the interaction between USP7 and endogenously expressed proteins, a Streptavidin-binding peptide-FLAG (SFB)-tagged USP7 construct^57^ (Addgene plasmid #99393) was transfected into HEK293T cells via PEI transfection. The lysate containing SFB-USP7 was generated with TNE lysis buffer as described above and incubated with PBS-washed Dynabeads^TM^ MyOne^TM^ Streptavidin T1 beads (Invitrogen #65601) for 1 hour at 4 °C with rotation. Next, the beads were washed 3 times with TNE lysis buffer and 4 times with cold PBS, followed by LDS sample buffer elution. The eluted proteins were analyzed by western blot along with 2% input lysates in the same blot.

#### Tandem ubiquitin binding entities (TUBE) assay

The TUBE assay was performed according to Tsao et al 2021 with modification^58^. Briefly, primary cortical neurons from E15 mouse embryos were transduced with Cre wild-type (WT) or catalytically dead (CD) lentivirus at DIV3 and treated with 10μM MG132 at DIV5. Neuron lysates were generated with TNE lysis buffer 16 hours after MG132 treatment. Cold TBST-washed pan-selective TUBE 2 beads (LifeSensors #UM-0402M) were incubated with neuron lysates overnight at 4°C with rotation. Next, the beads were washed 5 times with cold TBST and 2 times with cold PBS, followed by LDS sample buffer elution. The eluted proteins were analyzed by western blot along with 2% input lysates in the same blot.

#### *In vitro* deubiquitination assay

For *in vitro* ubiquitination assay, HEK293T cells were transfected with pcDNA3.1-FLAG-Ppil4 plasmid. After the treatment of 10µM proteasome inhibitor MG132 for 16 hours, the cells were harvested, lysed with 1% NP40 TNE lysis buffer, and the FLAG-Ppil4 was purified by immunoprecipitation with M2 anti-FLAG agarose beads (Sigma) as described above. 20μL PBS-washed M2 beads were added to 2.5mL lysate at 2μg/μL total protein concentration for 3 hours incubation at 4°C. The beads were washed 3 times with TNE lysis buffer, 2 times with high-salt PBS buffer (0.5M NaCl) to reduce non-specific binding, and then 3 times with regular PBS. Then we resuspended the washed beads with deubiquitination buffer (50mM Tris-HCl, 150mM NaCl, 5mM MgCl_2_ and 10mM DTT, final pH 7.3) and distributed 9µL beads for each deubiquitination reaction. Either 1µL 500nM recombinant His-USP7 (R&D system) or 1µL deubiquitination buffer as vehicle control was added to each reaction and incubated for 1 hour at 37 °C. LDS sample buffer was then added to stop the reaction and ubiquitinated Ppil4 was analyzed with western blot. Densitometric analysis was performed with FLAG-Ppil4 purified from 3 biological replicates of HEK293T lysates. Ubiquitinated Ppil4 was quantified as the fraction of Ppil4 integral fluorescent intensity above the molecular weight of the lowest full-length Ppil4 (∼65kD).

#### Diffusion-weighted *ex vivo* MRI and volumetric measurements

The skull of the fixed brain was removed before transferring to a 3-mL syringe containing fresh PBS. Diffusion-weighted MRI experiment was performed on 9.4-T Bruker scanner with a CryoProbe™. Fifteen sagittal slices, centered at the midline (slice 8), were acquired to cover the entire brain. Twenty-five-direction diffusion-weighted measurements were performed using a spin-echo diffusion-weighting sequence with the following parameters: repetition time (TR) = 1.5 s; echo time (TE) = 32 ms; field of view (FOV) = 20 × 20 mm2; matrix size = 192 × 192 (zero-filled to 384 × 384), slice thickness = 0.6 mm, max; b value = 3,000 s/mm2, Δ = 20 ms; δ = 5 ms; partial Fourier factor = 1.33; total scan time = 93.6 min. Diffusion-weighted data were converted to nifiti format followed by analysis using the lab-developed DBSI software package to perform DBSI multi-tensor analysis^59^. DBSI-derived fiber fraction was used to reflect apparent axonal^60^/dendritic^61^ density. DBSI-derived restricted and non-restricted fractions were used to estimate the extent of cellularity and vasogenic edema^61^. For volumetric measurement, segmentation was performed manually employing ITK-SNAP (version 3.8, www.itksnap.org)^62^. The Allen Mouse Brain Atlas was used as a reference to identify cortex and hippocampus. The three parasagittal slices at each side of midline (slices 4 −6 and 10 −12) were used to segment cortex and hippocampus.

#### Adeno-associated virus (AAV) injection

The AAV transfer plasmid pAAV-hSyn1-EGFP-P2A-EGFPf-WPRE-HGHpA was a gift from Dr. Guoping Feng (Addgene # 74513). The AAV PHP.eB was packaged at Hope Center Viral Vector Core of Washington University School of Medicine and diluted to 1.5 x 10^7^ vg/μL with 5% D-sorbitol in PBS. For intraventricular injection, P0/1 pups were anesthetized with hypothermia. 1uL of the diluted AAV was injected into each hemisphere (0.8-1mm away from the middle point between Lambda and Bregma, 2mm deep) of the pups. The injected mice were placed onto heat pad for body temperature recovery before returned to dam in the cage. We then perfused the injected mice at P18 and their brains were proceeded to cryosectioning and immunofluorescence of GFP for spine imaging and analysis.

#### Airyscan imaging and spine analysis

We used Zeiss 880 LSM2 confocal microscope with Airyscan hexagonal detector array and oil immersion to image z-stacks of dendritic spines at near-super resolution. For *in vivo* spine analysis, coronal sections at 100um thickness were float-stained for GFP. Based on Allen mouse reference atlas, motor cortices (M1 and M2) were imaged and 3-8 pyramidal neurons per section, 4 consecutive sections per animal, 3 animals per genotype were analyzed. The analysis was restricted to secondary apical dendritic segment and secondary or tertiary basal dendritic segment, starting from less than 40um distance to soma and for over 30um length. Representative images of dendritic spines of four shapes (mushroom, thin, stubby and filopodial) were pre-selected before quantification and used as references to compare with all other dendritic spines to call their shape in a blinded manner. For *in vitro* spine analysis, primary cortical neurons were seeded onto #1.5 coverslips. The neurons were transduced with lentivirus of MISSION TRC1-shRNA (Sigma) knocking down candidate substrates at DIV4, sparsely transfected with EGFP-expressing plasmid at DIV10, fixed at DIV18, and proceeded to immunocytochemistry of GFP and PSD95. Dendritic spines of 4-12 neurons were imaged per well, 3 wells per group, from 2 batches of neuron culture. The analysis was restricted to primary, 2^nd^ and 3^rd^ dendrites starting from less than 40μm distance to soma and for over 30μm length. The spines and PSD95 puncta were counted with Zeiss ZEN software in three dimensional view on a blinded basis. We used Imaris (Oxford Instruments) filament tracing to measure dendrite length in three dimensional view and divide counted number of spines and PSD95 puncta to get spine and PSD95 puncta density, respectively.

#### Dendrite tracing and morphological analysis

To assess dendrite morphology, primary cortical neurons from *Usp7* ^fl/fl^; *Trp53* ^+/−^ embryos were seeded onto #1 coverslips and transduced with Cre lentivirus at DIV3. The neurons was sparsely transfected with EGFP-expressing plasmid at DIV9, fixed at DIV11 for immunocytochemistry of GFP. The z-stacks of GFP signal were imaged with Zeiss 880 LSM2 confocal microscope and regular 20x objective. Based on the GFP signal, we used Imaris (Oxford Instruments) to model soma shape and trace dendrite morphology with filament-Autopath algorithm. Physically crossing dendrites were manually separated. Statistics of soma volumes, dendrite lengths and Sholl intersections were all determined and exported with Imaris filament. Complete dendrites of 9-15 neurons per well, 3 wells per group, from 2 batches of neuron culture were imaged and analyzed. All image acquisition and analysis were performed on a blinded basis.

#### MTS assay

Viability of the primary cortical neurons was assessed according to our previous publication^63^. Briefly, MTS reagent (Promega) was added to neurons 1:5 ratio to medium volume at DIV11 according to manufacturer’s instructions. The treated neurons were incubated at 37 °C with 5% CO_2_ for 2 hours before measurement of absorbance at 490nm. The measured MTS absorbance was subtracted with background signal in empty wells and normalized to control group (Cre dead or untreated neurons).

### Quantification and statistical analysis Gene

#### set enrichment analysis (GSEA)

GSEA software 4.3.2^64,65^ was used to identify the significantly differential enrichment of annotated gene sets between proteins in cortices of USP7 WT and Usp7 cKO mice. Gene sets were from M5 (gene ontology) of Molecular signatures database. NES values were calculated and gene sets with p-value < 0.05 were loaded into Cytoscape 3.9.1 for network construction. The enrichment map was constructed with yfiles Organic Layout, q-value < 0.01 as node cutoff, and similarity < 0.5 as edge cutoff. All gene sets shown in the enrichment map were enriched in USP7 WT mice and down-regulated in Usp7 cKO mice, as indicated by the green color (color scheme: PiYG-3).

#### Gene ontology analysis

GOrilla^22^ (http://cbl-gorilla.cs.technion.ac.il) was used to identify significantly enriched gene ontology terms in ranked gene list. For *in vivo* TMT protein abundance, n=131 proteins down-regulated in Usp7 cKO mice (Fold Change < 0.75) were ranked by Fold Change and loaded to GOrilla with a background of total proteins detected in all mice. 3 uncharacterized proteins without gene name were not detected by GOrilla. No GO term was significantly enriched in *Usp7* cKO up-regulated proteins.

#### SynGO analysis

For SynGO analysis, unranked down-regulated proteins in *Usp7* cKO; *Trp53*^+/−^ cortex in TMT-mass spectrometry experiment were identified with cutoff of Fold Change < 0.75 and 2-tail unpaired t-test p-value < 0.01. All proteins detected in all mice were used as background. The SynGO enrichment analysis was performed with the SynGO (v1.2) online tool for both cellular components and biological processes (https://syngoportal.org)^66^.

#### STRING network analysis

The String network was built with StringApp 2.0.1^67^ in Cytoscape 3.9.1. To ensure specificity, we calculated the enrichment percentage of the spectral counts in USP7 TRAF WT group, and only interactors with over 95% enrichment were proceeded for the following analysis. Top 50 interactors based on spectral counts normalized to protein length were converted into homologous mouse proteins and loaded into StringApp to create the association map. Two uncharacterized rat proteins were not included. Full STRING network with edges of confidence over 0.4 was built. Enriched molecular components (Gene ontology) were assigned to subunits of the complex in the association map. FDR of each enriched molecular components was calculated using the Benjamini-Hochberg procedure.

#### Disease gene enrichment analysis

Interactors of USP7 TRAF domain used in the disease gene enrichment analysis are the detected proteins in all 3 biological replicates of IP-mass spectrometry experiment. The interactors were filtered with 95% enrichment in USP7 TRAF WT group, as described in the String network analysis. All proteins in primary neuron culture detected by TMT-mass spectometry were used as background. Curated pathogenic gene sets for autism, intellectual disability (ID), Epilepsy, ADHD, schizophrenia, bipolar disorder and cerebral palsy were from Geisinger developmental brain disorder gene database (https://dbd.geisingeradmi.org/)^24^. High-confidence FMRP targets were from FMRP enriched genes in HITS-CLIP^68^. USP7 TRAF interactors, FMRP targets, and the background proteins were all converted into human orthologs entrez ID with biomaRt R package before the analysis. p-values of the disease gene set enrichment were calculated with one-sided hypergeometric test, and then adjusted by Benjamini-Hochberg procedure.

#### Principal component analysis (PCA)

PCA of all 16 samples of the *in vitro* TMT experiment was performed with function prcom() of R. All 8304 detected proteins were loaded into the analysis and principal component 1 and 2 were mapped with variance percentage.

#### Selection of candidate substrates

To filter out proteins with little to no expression in glutamatergic neurons, we used Allen Institute Smart-Seq database^69^ to calculate the total CPM of the corresponding gene in the clade of glutamatergic neuron clusters. Proteins with total CPM level in glutamatergic neurons below that of the astrocyte marker aldolase C were excluded from the following analyses. To select candidate substrates of USP7, we first calculated the Euclidean distance of each protein to USP7 over all 16 samples and sorted out the 25 proteins closest to USP7 as majority of the candidate substrates. Because the sorting based only on Euclidean distance overlooks the proteins with differential abundances between the two time points (DIV6 and DIV8), a 2×2 two-way ANOVA was also applied to all proteins to calculate the p-values of Cre and Cre-time interaction effect on the abundance of each protein. Proteins down-regulated upon wild-type Cre with p-values < 10^−7^ were added to the candidate substrate list.

#### Weighted gene correlation network analysis (WGCNA)

The R package WGCNA^70^ was used to construct a co-expression network for all the proteins with significant Cre effect (n=2307) or Cre-time interaction effect (n=320) (p<0.05 by two-way ANOVA). All 16 TMT samples were used to calculate Pearson’s correlation matrices representing co-expression similarity of proteins, and different Cre type (Cre WT and Cre dead) and time points (DIV6 and DIV8) were loaded as traits. The weighted adjacency matrix was created using Pearson correlation coefficient test of all differentially expressed proteins and transformed into a topological overlap measure (TOM) matrix and average linkage hierarchical clustering of the TOM matrix was used to construct a clustering dendrogram of the genes with thresholding power 12. 10 Modules in different colors were generated with minimal gene module size of 30 proteins and minimal height to merge modules of 0.15. GO term enrichment analyses for the modules were performed by DAVID 2021 Functional Annotation Tool^71,72^ with a background of all module-assigned proteins. For module-trait relationship, Cre WT, Cre dead, DIV6 and DIV8 were assigned as 1, 0, 6 and 8 respectively. The correlation coefficient and correlation p-value of the first principal component of the module gene expression profile (eigengene) and traits were calculated to identify trait-correlated modules.

#### Statistics

All statistical analyses were performed using IBM SPSS Statistic software (v.26), GraphPad Prism (v.9) and R. For experiments to compare only two groups, student t-test was used. Analysis of variance (ANOVA), (one-way, two-way, factorial, repeated measures) were used analyze data with multiple groups. With a statistically significant interaction between main factors, simple main effects were calculated to provide clarification of statistically significant between-group and within-group differences. Where appropriate, the Huynh-Feldt adjustment was used to protect against violations of sphericity. For multiple pairwise comparisons and post-hoc tests, Tukey correction was applied to compare every two means, and Bonferroni correction was applied to compare selected means when independence cannot be assumed. For three groups with non-normal distributions (sensorimotor battery), the nonparametric Kruskal-Wallis test was used followed by post hoc test with the Dunn correction to compare every mean to a control mean. All data in bar graphs were presented as mean ± SEM and probability value for all analyses was *p < 0.05, **p < 0.01, ***p < 0.001, ****p < 0.0001, unless otherwise stated.

---

**Figure S1.**
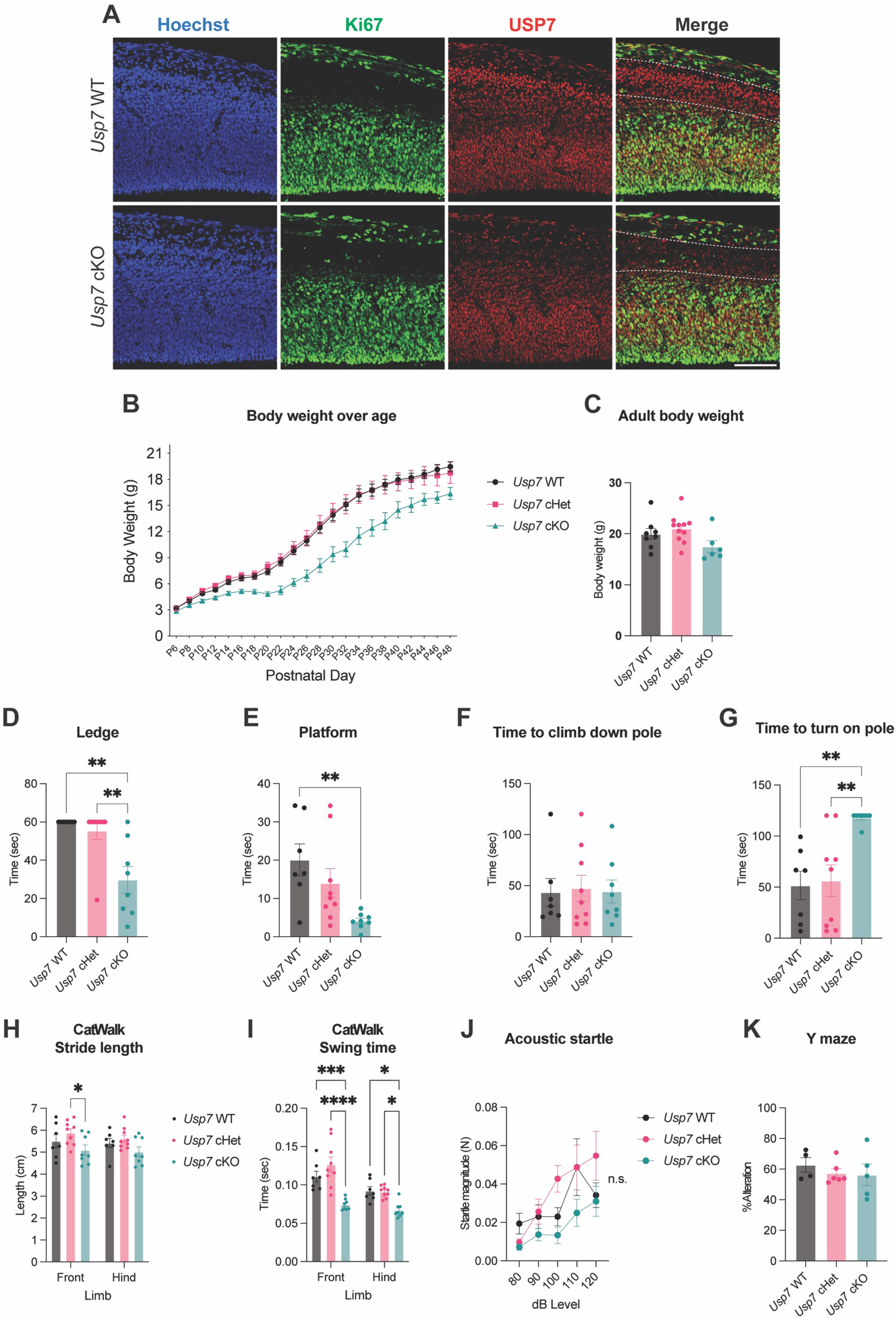
Additional phenotypic characterization of *Usp7* cKO mice. Related to Figure 1 and Table 1. (A) Immunofluorescence of USP7 in cerebral cortex at embryonic day 14 (E14). USP7 is localized to both Ki-67+ ventricular-subventricular zone (V-SVZ) and cortical plate (CP) in USP7 WT cortex. But in *Usp7* cKO cortex, USP7 is depleted only in post-mitotic cortical plate (dashed outline). Scale bar 100µm. (B) Body weight curve from P6 to P48. *Usp7* cKO mice (n = 7) exhibited body weight growth delay, compared to *Usp7* WT (n = 10) and cHet mice (n = 7). (C) Body weight of adult mice. Number of mice = 8, 11 & 9 for *Usp7* WT, cHet & cKO, respectively. No significant genotype effect by one-way ANOVA. (D) Motor coordination test 1: Ledge test. *Usp7* cKO mice (n = 8) showed shorter latency to fall from ledge than *Usp7* WT (n = 7) and cHet mice (n = 9). **p<0.01 by Dunn’s multiple comparison test. (E) Motor coordination test 2: Platform test. *Usp7* cKO mice showed shorter latency to fall from platform than *Usp7* WT mice. Numbers of mice are the same as in (D). **p<0.01 by Dunn’s multiple comparison test. (F) Motor coordination test 3: Pole test, time to climb down the pole. Numbers of mice are the same as in (D). No significant genotype effect by one-way ANOVA. (G) Motor coordination test 4: Pole test, time to turn at the tip of the pole. *Usp7* cKO mice spent more time in turning than *Usp7* WT and cHet mice. Numbers of mice are the same as in (D). *p<0.05, **p<0.01 by Dunn’s multiple comparison test. (H) Catwalk 3: Stride length. *Usp7* cKO mice showed slightly shorter average stride length than control. Numbers of mice are the same as in (D). *p<0.05 by Tukey’s multiple comparison test. (I) Catwalk 4: Swing time. *Usp7* cKO mice used less time swinging front and hind legs for each step, compared to *Usp7* WT and cHet mice. Numbers of mice are the same as in (D). *p<0.05, ***p<0.001, ****p<0.0001 by Tukey’s multiple comparison test. (J) Hearing test: Magnitude of response to acoustic startle at different dB level. Number of mice = 5, 8 & 6 for *Usp7* WT, *Usp7* cHet & *Usp7* cKO. No significant genotype effect by two-way ANOVA. (K) Y maze: Alternative rate in choosing arm to enter. Number of mice = 4, 6 & 5 for *Usp7* WT, *Usp7* cHet & *Usp7* cKO. No significant genotype effect by one-way ANOVA. Data are presented as mean ± SEM.

**Figure S2.**
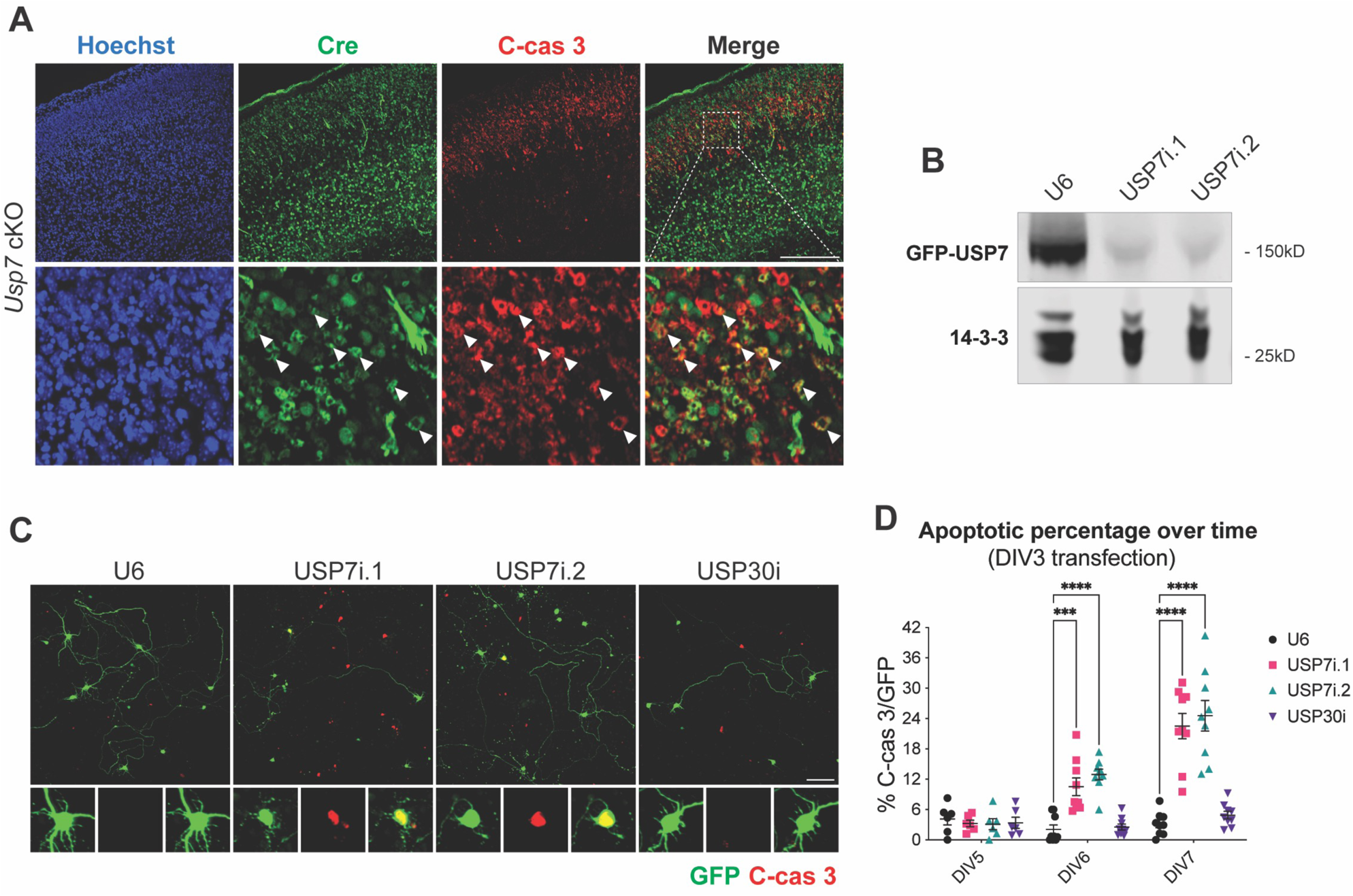
Loss of USP7 in glutamatergic neurons induces apoptosis in a cell autonomous manner. Related to Figure 2. (A) Double immunofluorescence of cleaved caspase 3 (c-cas 3) and Cre. An expanded region shows overlap (arrows) of cleaved caspase 3 and Cre, supporting that the apoptotic cells are glutamatergic neurons. Scale bar 200µm. (B) Western blot validation of USP7 RNAi in HEK293Tcells. Both USP7i.1 and USP7i.2 diminished expression of GFP-USP7. 14-3-3 is loading control. (C) Immunocytochemistry of c-cas 3 in primary cortical neurons dissociated from wild-type E14 cortices and transfected with a GFP plasmid. Scale bar 100µm. Cells were fixed at DIV5, 6 and 7. At the bottom are close-up of neurons with and without c-cas 3 signal. (D) Quantification of apoptotic percentage, defined as the percentage of c-cas 3 and GFP double positive neurons among all GFP positive neurons. Cortical neurons were transfected with USP7 shRNA plasmids at 3 days *in vitro* (DIV3) and analyzed at DIV5, 6 and 7. Data were collected from three independent batches of neuron culture, three wells per condition within each batch. USP7 knockdown by two independent shRNA led to a progressive elevation in apoptotic percentage. Empty vector U6 and knockdown of an irrelevant deubiquitinase, USP30, served as negative controls. ***p<0.001, ****p<0.0001 by Tukey’s multiple comparison test. Data are presented as mean ± SEM.

**Figure S3.**
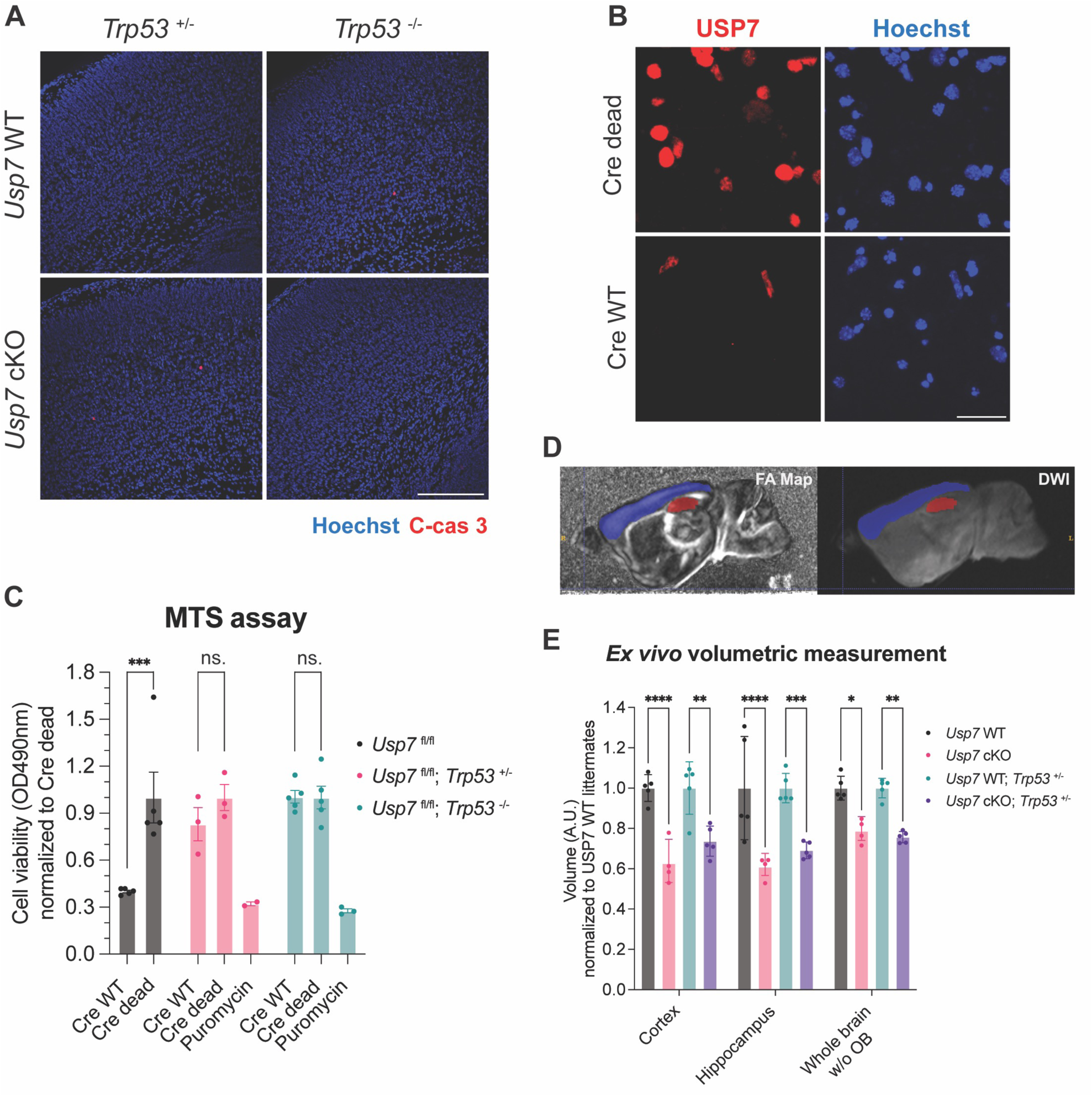
Deletion of p53 rescues USP7 loss-induced apoptosis but minimally restores the brain volume reduction in *Usp7* cKO mice. Related to Figure 2 and Table 1. (A) Immunofluorescence of c-cas 3 in the cerebral cortex of *Usp7* cKO mice at P0 with deletion of one or both allele of *Trp53*. C-cas 3 signal remained sparse with either *Trp53* heterozygosity or homozygosity. Scale bar 200µm. (B) Immunofluorescence of USP7 in primary cortical neurons dissociated from *Usp7*^fl/fl^; *Trp53*^+/−^ embryos. Neurons transduced with Cre WT showed loss of USP7 protein at DIV14. Scale bar 25µm. (C) MTS assay of *Usp7*^fl/fl^ cortical neurons transduced by lentiviral Cre with different *Trp53* genotypes. USP7 knockout in *Trp53*^+/+^ neurons (n = 5 wells) strongly reduced cell viability, indicating of neuronal death. No significant viability decrease in *Trp53*^+/−^ (n = 3 wells) and *Trp53*^−/−^ neurons (n = 5 wells). Puromycin (2µg/mL, 24hrs) treated neurons serve as positive control. ***p<0.001 by Šídák’s multiple comparison test between Cre WT and Cre dead. (D) Representative segmentation on FA map and DWI for volumetric measurement with diffusion-weighted *ex vivo* MRI. Cerbral cortex and hippocampus are shown in blue and red, respectively. (E) Volumetric measurement with *ex vivo* MRI. *Usp7* cKO brains (P30-32) are smaller than *Usp7 WT*. Deletion of p53 minimally stored the volume of cortex and hippocampus. OB = olfactory bulbs. Number of mice = 5, 4, 5 & 5 for *Usp7* WT, *Usp7* cKO, *Usp7* WT; *Trp53* ^+/−^ & *Usp7* cKO; *Trp53* ^+/−^. *p<0.05, **p<0.01, ***p<0.001, ****p<0.0001 by Tukey’s multiple comparison test. Data are presented as mean ± SEM.

**Figure S4.**
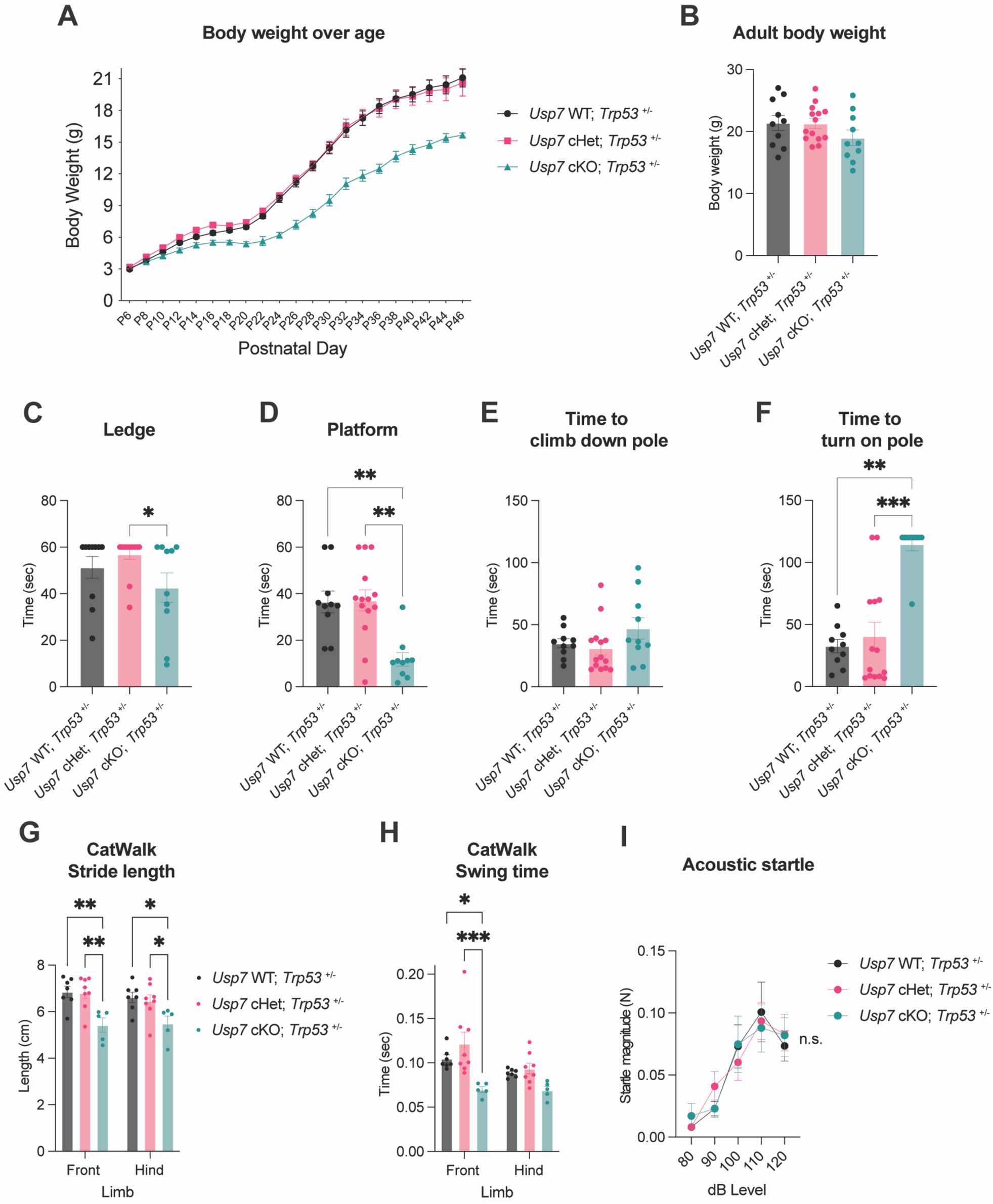
Additional phenotypic characterization of *Usp7* cKO; *Trp53*^+/−^ mice. Related to Figure 2 and Table 1. (A) Body weight curve from P6 to P48. *Usp7* cKO; *Trp53*^+/−^ mice (n = 6) exhibited body weight growth delay, compared to *Usp7* WT; *Trp53*^+/−^ (n = 10) and *Usp7* cHet; *Trp53*^+/−^ mice (n = 5). (B) Body weight of adult mice. Number of mice = 10, 14 & 10 for *Usp7* WT, cHet & cKO; *Trp53* ^+/−^, respectively. No significant genotype effect by one-way ANOVA. (C) Motor coordination test 1: Ledge test. *Usp7* cKO; *Trp53*+/− mice showed slightly shorter latency to fall from ledge than control mice. Numbers of mice are the same as in (B). *p<0.05 by Dunn’s multiple comparison test. (D) Motor coordination test 2: Platform test. *Usp7* cKO; *Trp53*^+/−^ mice showed shorter latency to fall from platform than control mice. Numbers of mice are the same as in (B). **p<0.01 by Dunn’s multiple comparison test. (E) Motor coordination test 3: Pole test, time to climb down the pole. Numbers of mice are the same as in (B). No significant genotype effect by one-way ANOVA. (F) Motor coordination test 4: Pole test, time to turn at the tip of the pole. *Usp7* cKO; *Trp53*^+/−^ mice spent more time in turning than control mice. Numbers of mice are the same as in (B). **p<0.01, ***p<0.001 by Dunn’s multiple comparison test. (G) Catwalk 3: Stride length. *Usp7* cKO; *Trp53*^+/−^ mice showed shorter average stride length than control. Number of mice = 7, 8 & 5 for *Usp7* WT, cHet & cKO; *Trp53* ^+/−^. *p<0.05, **p<0.01 by Tukey’s multiple comparison test. (H) Catwalk 4: Swing time. *Usp7* cKO; *Trp53*^+/−^ mice used less time swinging front and hind legs for each step, compared to control mice. Numbers of mice are the same as in (G). *p<0.05, ***p<0.001 by Tukey’s multiple comparison test. (I) Hearing test: Magnitude of response to acoustic startle at different dB level. Number of mice = 10, 14 & 10 for *Usp7* WT, cHet & cKO; *Trp53* ^+/−^. No significant genotype effect by two-way ANOVA. Data are presented as mean ± SEM.

**Figure S5.**
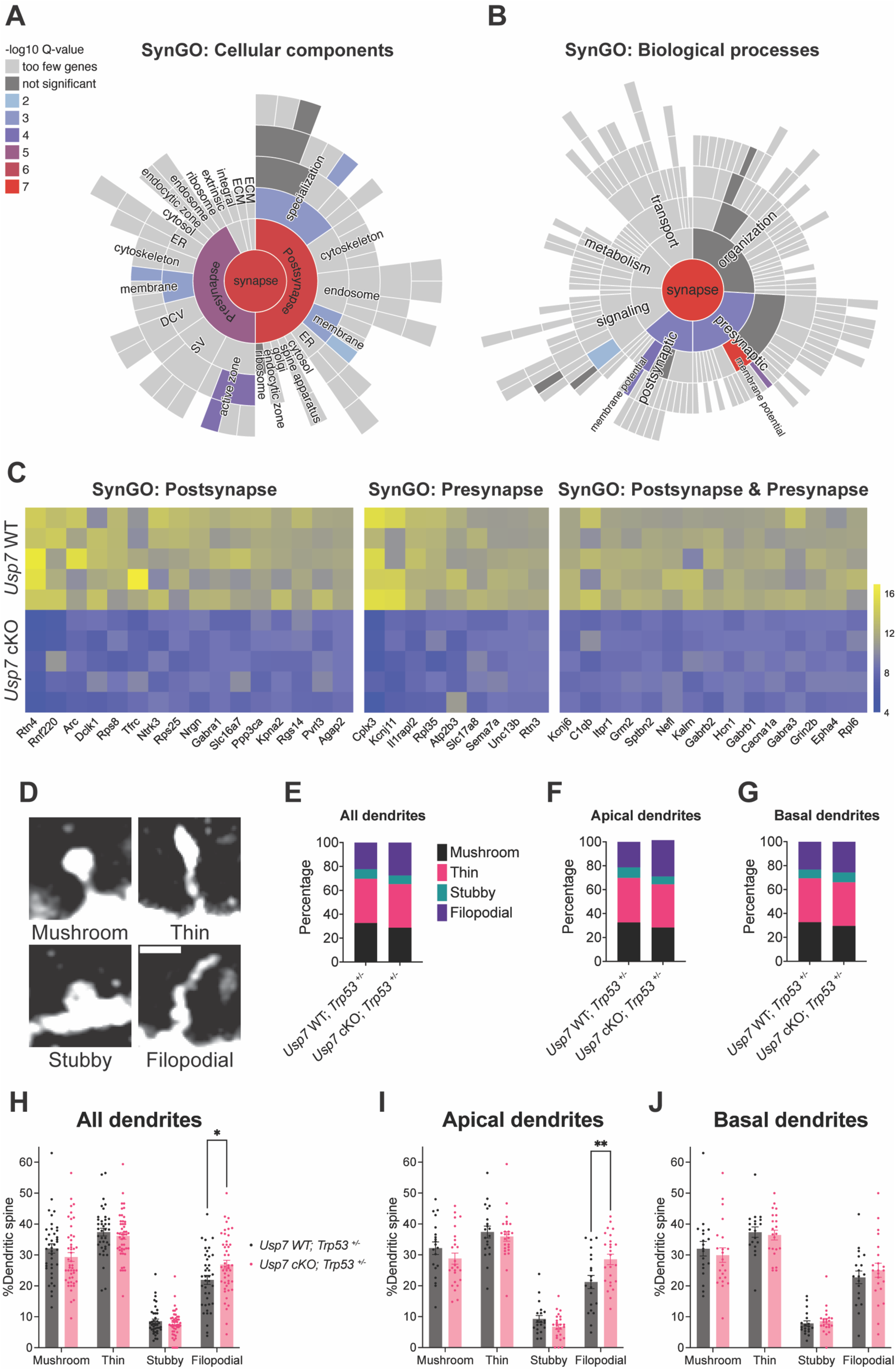
Additional analyses of reduced synaptic proteins and dendritic spine morphology in *Usp7* cKO mice. Related to Figure 3 (A&B) Sunburst plot of SynGO terms enriched in down-regulated proteins of the TMT proteome of *Usp7* cKO cortex. Cellular component SynGO terms in (A), biological process SynGO terms in (B). (C) TMT relative abundances of down-regulated synaptic proteins in *Usp7* cKO cortex. The individual proteins within SynGO: Postsynapse, SynGO: Presynapse and in both were segregated into different panels. (D) Representative GFP micrographs of dendritic spines of different shapes, including mushroom, thin, stubby and filopodial spines. Scale bar 1µm. (E-J) Percentages of dendritic spines of different shapes in *Usp7* cKO and WT cortex. (E-G) shows total percentages of dendritic spines of different shapes. (H-J) shows by neuron percentages of dendritic spines of different shapes. There was an increase in the proportion of filopodial dendritic spines in *Usp7* cKO pyramidal neurons (n = 25 & 22 neurons for apical dendrites & basal dendrites from 3 mice), compared to *Usp7* WT pyramidal neurons (n = 20 & 19 neurons for apical dendrites & basal dendrites from 3 mice). *p<0.05, **p<0.01 by Šídák’s multiple comparison test. Data are presented as mean ± SEM.

**Figure S6.**
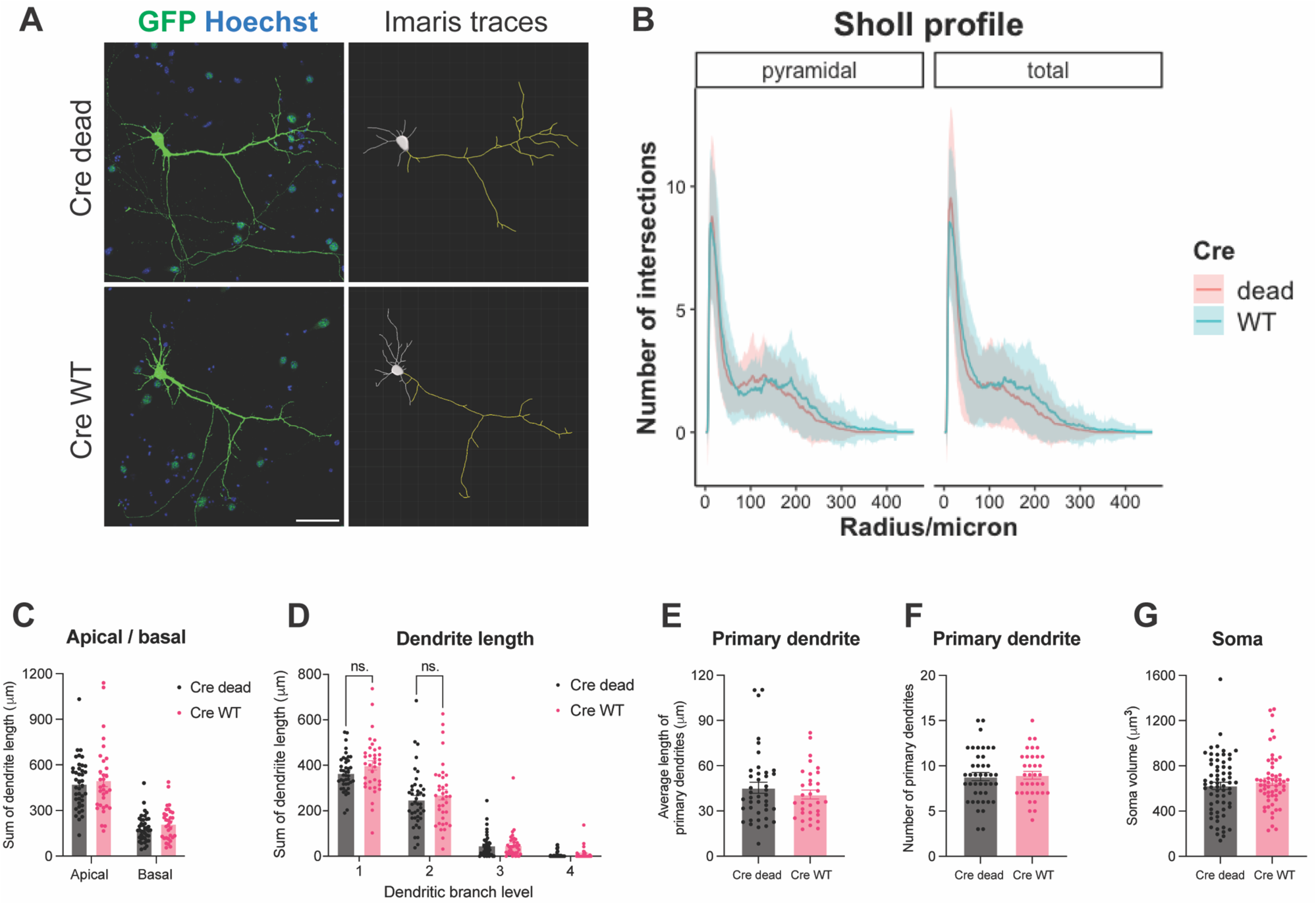
USP7 loss does not cause significant change in dendritic morphology in primary cortical neurons with Trp53 heterzygosity. Related to Figure 3. (A) Representative DIV11 images of cortical pyramidal neurons and their Imaris dendritic traces. For the Imaris dendritic traces, apical dendrties are highlighted in yellow. The cortical neurons were dissociated from *Usp7*^fl/fl^; *Trp53*^+/−^ E15 embryos and transduced with Cre WT or Cre dead lentivirus at DIV3. (B) Sholl profile of pyramidal neurons (n = 40 & 32 for Cre dead & WT) and all neurons (n = 55 & 44 for Cre dead & WT). Shaded area represents standard deviation. Profiles of neurons transduced with Cre dead and Cre WT are colored in red and green, respectively. (C) Sum of dendrite length in pyramidal neurons, apical dendrites vs. basal dendrites (n = 40 & 32 for Cre dead & WT). No significant Cre effect by two-way ANOVA. (D) Sum of dendrite length in pyramidal neurons, across 1-4 branch levels. Numbers of neurons are the same as in (C). No significant Cre effect by two-way ANOVA. (E) Average length of primary dendrites in pyramidal neurons. Numbers of neurons are the same as in (C). No significance by 2-tail unpaired t-test. (F) Number of primary dendrites in pyramidal neurons. Numbers of neurons are the same as in (C). No significance by 2-tail unpaired t-test. (G) Soma 3D-volumes of pyramidal neurons (n = 65 & 57 for Cre dead & WT). No significance by 2-tail unpaired t-test. Data are presented as mean ± SEM.

**Figure S7.**
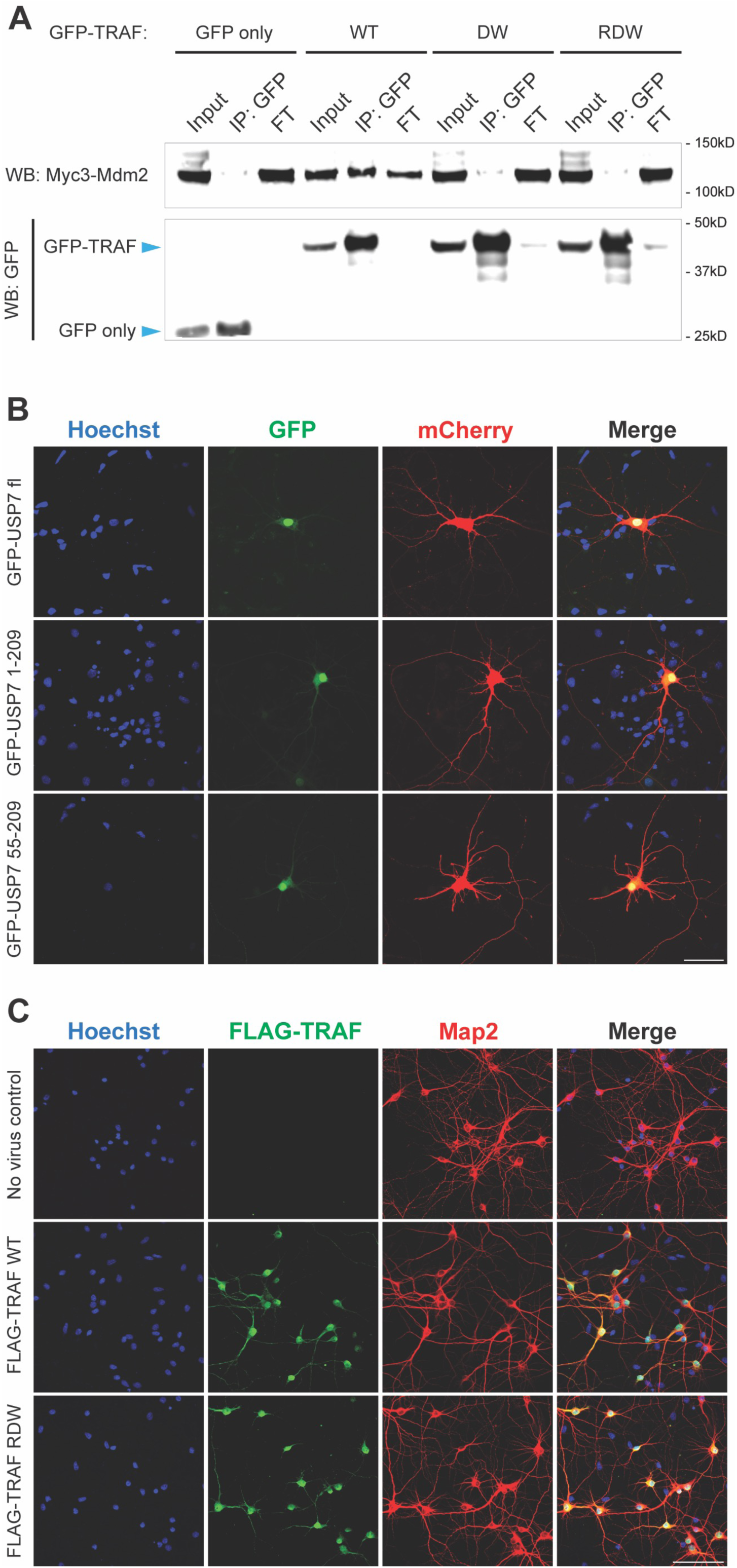
Validation of USP7 TRAF domain expression and interaction for IP-MS. Related to Figure 4. (A) Immunoprecipitation of GFP-TRAF domain followed by western blot of myc3-Mdm2 and GFP. Lysates were prepared from HEK293T cells transfected with myc3-Mdm2 and GFP-tagged USP7-TRAF domain. DW double and RDW triple mutations (All mutated into Alanine) on the USP7-TRAF domain disrupted the interaction with MDM2. FT as flow-through lysates. (B) Mostly nuclear localization of truncated GFP-USP7 in rat cortical neurons at DIV9, including USP7-TRAF domain (USP7 amino acid 55-209). Neurons were co-transfected with pCAG-mCherry with Ca phosphate. USP7 fl as USP7 full length. Scale bar 50µm. (C) Immunocytochemistry of FLAG showed mostly nuclear expression of FLAG-USP7 TRAF domain in rat cortical neurons with and without RDW mutations at DIV10. Immunocytochemistry of Map2 shows neuronal morphology. Scale bar 100µm.

**Figure S8.**
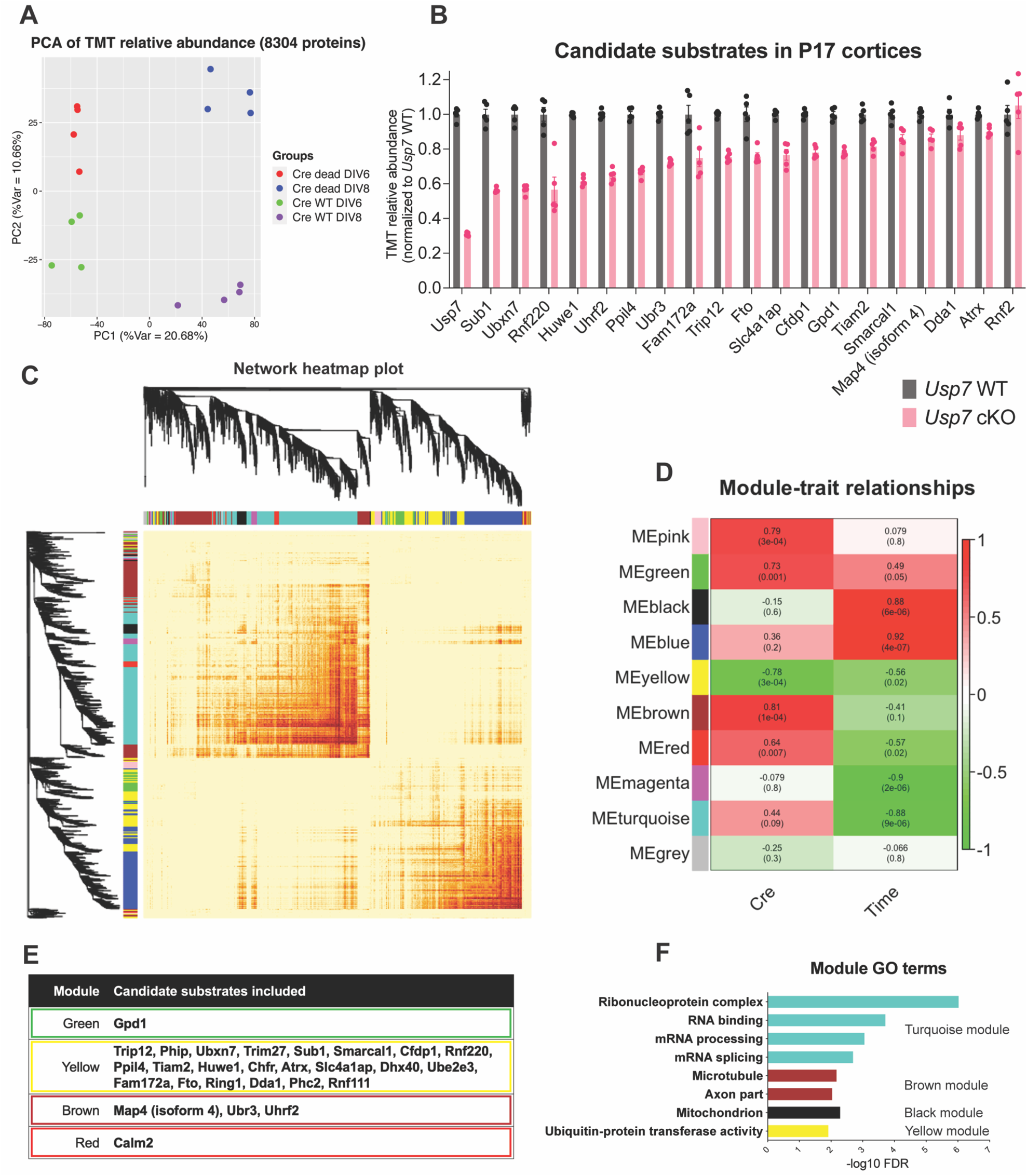
Additional analyses of the TMT proteome with acute USP7 loss in cortical neurons. Related to Figure 4. (A) PCA of 16 samples based on TMT abundance of all 8304 proteins detected by TMT-MS. Principal component 1 (PC1) separates DIV6 and DIV8 samples whereas principal component 2 (PC2) separates Cre WT and Cre dead samples. (B) TMT protein abundance of candidate substrates in previous *in vivo* TMT-MS (Figure 3A-D). Most candidate substrates showed various degrees of reduction in the cerebral cortex of *Usp7* cKO mice at P17. n = 5 for both *Usp7* WT; *Trp53* ^+/−^ and *Usp7* cKO; *Trp53* ^+/−^ mice. (C) Dendrogram and heatmap of TMT protein adjacencies by weighted gene co-expression network analysis (WGCNA). WGCNA segragates 2627 differentially expressed proteins (Cre effect p<0.05 or Cre time interaction effect p<0.05 by two-way ANOVA) into 10 modules in different colors. (D) Module-trait relationships. Correlations and p-values between module eigengene (ME) and Cre (Cre WT as 1, Cre dead as 0) or time (DIV6 or DIV8) are denoted and color-coded according to the color legend. (E) Membership of candidate substrates in each module. (F) GO annotation for each module. Turquiose module is enriched in ribonucleoprotein complex and splicing-related terms. Brown module is enriched in microtubule and axon-related terms. Black module is enriched in mitochondrion-related terms. Yellow module is enriched in E3 ligase-related terms. Data are presented as mean ± SEM.

**Figure S9.**
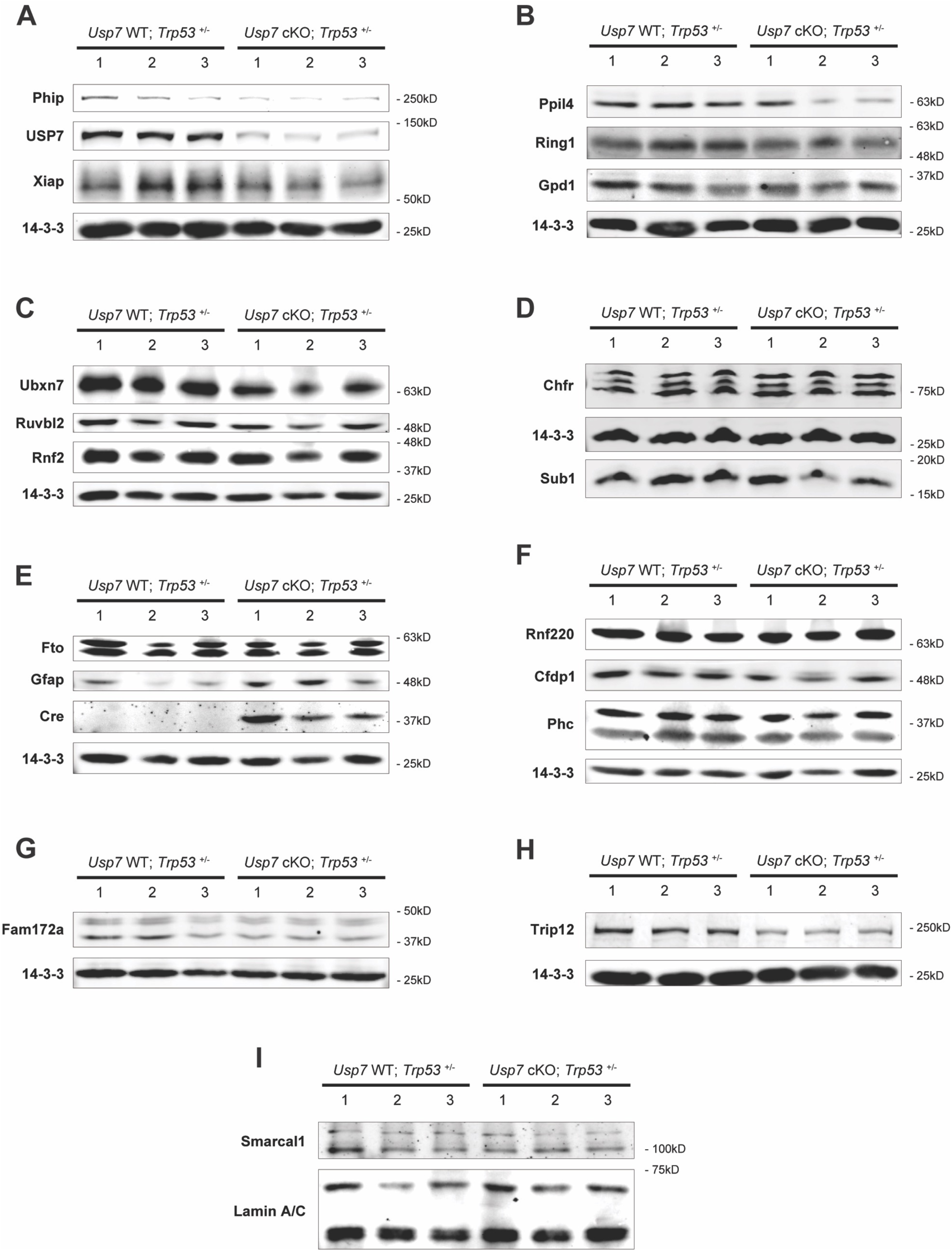
Western blot of candidate substrates in the cerebral cortex of P0 *Usp7* cKO; *Trp53*^+/−^ mice. Related to Figure 4(H). Three biological replicates of each genotype were denoted in numbers. (A) Phip, USP7 and Xiap. (B) Ppil4, Ring1 and Gpd1. (C) Ubxn7, Ruvbl2 and Rnf2. (D) Chfr and Sub1. (E) Fto, Gfap and Cre. (F) Rnf220, Cfdp1 and Phc. (G) Fam172a. (H) Trip12. (I) Smarcal1. 14-3-3 and Lamin A/C are loading controls.

**Figure S10.**
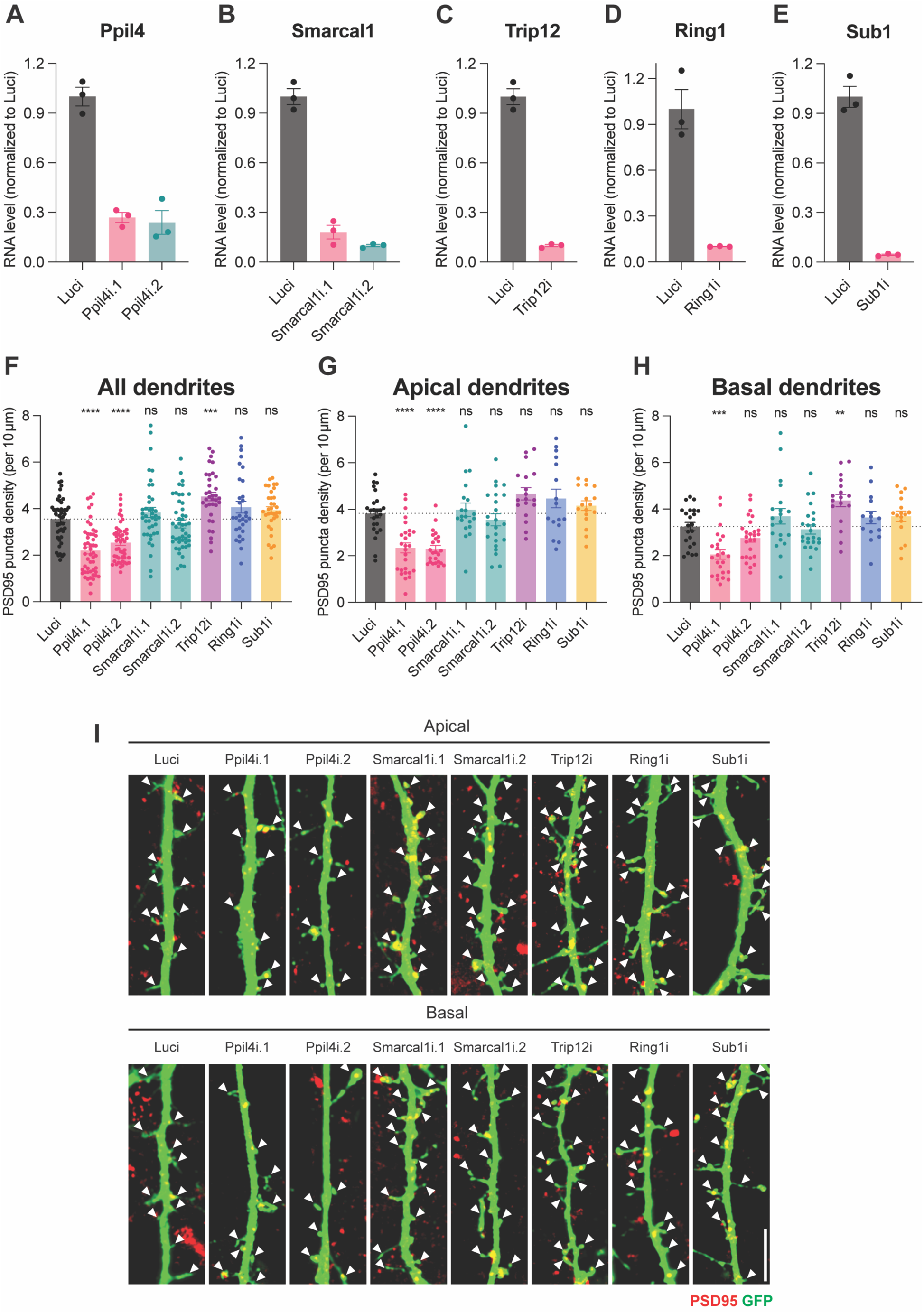
Additional characterization of the effect of candidate substrate knockdown on dendritic spine morphogenesis. Related to Figure 5 (A-E) RT-qPCR validation of candidate substrates RNAi (Ppil4, Smarcal1, Trip12, Ring1 and Sub1) in primary cortical neurons. Neurons were transduced with shRNA lentivirus at DIV4, lysed and harvested at DIV7. n = 3 biological replicates for all groups (F, G, H) Density of PSD95 puncta on apical, basal and all dendrites of pyramidal-shaped neurons in (Figure 5H). Two independent lentiviral shRNA knocking down Ppil4 reduced PSD95 puncta density compared with control shRNA targeting Luciferase (Luci). Number of neurons (apical dendrties) = 24, 27, 23, 20, 23, 18, 16 & 17 for Luci, Ppil4i.1, Ppil4i.2, Smarcal1i.1, Smarcal1i.2, Trip12i, Ring1i & Sub1i. Number of neurons (basal dendrties) = 22, 24, 26, 20, 24, 17, 15 & 16 for Luci, Ppil4i.1, Ppil4i.2, Smarcal1i.1, Smarcal1i.2, Trip12i, Ring1i & Sub1i. **p<0.01, ***p<0.001, ****p<0.0001 by Dunnett’s multiple comparison test (compared to Luci). (I) Representative images show the effect of knockdown of candidate substrates on dendritic spines of cortical neurons at DIV18. PSD95 puncta decorating the dendritic spines were immunostained along with GFP. Arrows show dendritic spines. Scale bar 5µm. Data are presented as mean ± SEM.

## Other Supplemental Materials

Table S1. GSEA statistic report of TMT proteomics of *Usp7* cKO and control cortex.

Table S2. Oligonucleotides used in this study.

Video S1 Abnormal gait of *Usp7* cKO mouse. Mouse age P33.

Video S2 *Usp7* cKO vs WT mouse CatWalk with 10x slow speed.

Video S3 Aggressive biting in *Usp7* cKO mouse. Mouse age P33.

Video S4 *Usp7* cKO; *Trp53*^+/−^ vs *Usp7* WT; *Trp53*^+/−^ mouse CatWalk with 10x slow speed.

