## Supplementary material for "The Hao-Fountain syndrome protein USP7 regulates neuronal connectivity in the brain via a novel p53-independent ubiquitin signaling pathway": Table S2

Table S5. Oligonucleotides used in this study.

| **Oligonucleotides** | **Sequence (5’-3’)** | **Source** |
| --- | --- | --- |
| *Cre* genotyping primer F | GCTAAACATGCTTCATCGTCGG | Yamada et al. 2014 |
| *Cre* genotyping primer R | GATCTCCGGTATTGAAACTCCAGC | Yamada et al. 2014 |
| *Usp7* floxed genotyping primer #1 | GAAGCATACTGGCTATGTCGGCTT | Kon et al. 2011 |
| *Usp7* floxed genotyping primer #2 | GCCTGTCAGCTACAAGAGAGCCAG | Kon et al. 2011 |
| *Usp7* floxed genotyping primer #3 | CATCAGTCAGGTACATAATATAAC | Kon et al. 2011 |
| *Tp53* knockout genotyping primer #1 | TGGATGGTGGTATACTCAGAGC | The Jackson Laboratory |
| *Tp53* knockout genotyping primer #2 | CAGCCTCTGTTCCACATACACT | The Jackson Laboratory |
| *Tp53* knockout genotyping primer #3 | AGGCTTAGAGGTGCAAGCTG | The Jackson Laboratory |
| Gapdh RT-qPCR primer F | TGCTGGTGCTGAGTATGTCG | Majidi et al. 2019 |
| Gapdh RT-qPCR primer R | GCATGTCAGATCCACAACGG | Majidi et al. 2019 |
| USP7 RT-qPCR primer F | CATGGAGTTGCGTGGGACTC | This paper |
| USP7 RT-qPCR primer R | CGCCCTCTGTTGGCATCATG | This paper |
| Ppil4 RT-qPCR primer F | AAAGCCCAAACAGGATGCAAAA | This paper |
| Ppil4 RT-qPCR primer R | GAGCAGTGGCGTGTTTTCTTT | This paper |
| Sub1 RT-qPCR primer F | TCAGTCCGGAGTAGGCGAG | This paper |
| Sub1 RT-qPCR primer R | TGATTTAGGCATTGCTCCGC | This paper |
| Ubxn7 RT-qPCR primer F | AAAAGGAGAAGGCCAGCACG | This paper |
| Ubxn7 RT-qPCR primer R | GCCACACTCTTTGGCTGTTTC | This paper |
| Trip12 RT-qPCR primer F | CGGAGCTGGAGGAAGCTTTT | This paper |
| Trip12 RT-qPCR primer R | TGCACAGTCTGGGTGTCTTT | This paper |
| Ring1 RT-qPCR primer F | TTGTGTTCCGGCCCCATC | This paper |
| Ring1 RT-qPCR primer R | CCAGTAGTCTTCACGTACCGAG | This paper |
| Fam172a RT-qPCR primer F | GTTGTAACTGGGTCTCCAGC | This paper |
| Fam172a RT-qPCR primer R | CTTGAAGATAGACGGGAAGCTC | This paper |
| Smarcal1 RT-qPCR primer F | TTCGACGGTGGCCGAGC | This paper |
| Smarcal1 RT-qPCR primer R | TTGGGAAGTTGGTAAAAGCTAGCGA |  |
| Cfdp1 RT-qPCR primer F | TTGTGGGCCAGCTTCCTTAAT | This paper |
| Cfdp1 RT-qPCR primer R | TTCAGTCTCCTCGGCTACCTTA | This paper |
| Chfr RT-qPCR primer F | CTGGAAGACACCAGCACCAA | This paper |
| Chfr RT-qPCR primer R | TCCCCGCTCTGTAAAGGGTA | This paper |
| USP7 RNAi construct #1 | CTAGAAATTGTGAGCTACAAA | Anckar et al. 2015 |
| USP7 RNAi construct #2 | GGATTATGTGGCAGTAGAACA | This paper |
| USP30 RNAi construct | TGATGGACTTCTACAAGTACC | Anckar et al. 2015 |
| Ppil4 RNAi construct #1 | CCAGCCAATTTGGTTCTGAAA | Sigma (TRC clone ID: TRCN0000101207) |
| Ppil4 RNAi construct #2 | CCGAGCCTACAAAGGAACAAT | Sigma (TRC clone ID: TRCN0000101208) |
| Smarcal1 RNAi construct #1 | GCCTATCCTAAAGGTTGCCAA | Sigma (TRC clone ID: TRCN0000109396) |
| Smarcal1 RNAi construct #2 | CGAGAGAAGTTTCTAGTGTTT | Sigma (TRC clone ID: TRCN0000109398) |
| Trip12 RNAi construct | GCAGGATAAATCCCAGACCAA | Sigma (TRC clone ID: TRCN0000039428) |
| Ring1 RNAi construct | CACTGACCTTGGAGCTTGTAA | Sigma (TRC clone ID: TRCN0000040561) |
| Sub1 RNAi construct | CCAGATTGGAAAGATGAGATA | Sigma (TRC clone ID: TRCN0000072049) |
| Luciferase RNAi construct | CAGAATCGTCGTATGCAGTGA | Genome Institute at Washington University (TRC clone ID: TRCN0000231714) |
